# SAMD1 suppresses epithelial-mesenchymal transition (EMT) pathways in pancreatic ductal adenocarcinoma

**DOI:** 10.1101/2023.08.14.553183

**Authors:** Clara Simon, Inka D. Brunke, Bastian Stielow, Ignasi Forné, Anna Mary Steitz, Merle Geller, Iris Rohner, Lisa M. Weber, Sabrina Fischer, Lea Marie Jeude, Andrea Nist, Thorsten Stiewe, Magdalena Huber, Malte Buchholz, Robert Liefke

## Abstract

SAMD1 (SAM-domain containing protein 1), a CpG island-binding protein, plays a pivotal role in the repression of its target genes. Despite its significant correlation with outcomes in various tumor types, the role of SAMD1 in cancer has remained largely unexplored. In this study we focused on pancreatic ductal adenocarcinoma (PDAC) and revealed that SAMD1 acts as a repressor of genes associated with epithelial-mesenchymal transition (EMT). Upon deletion of SAMD1 in PDAC cells, we observed significantly increased migration rates. SAMD1 exerts its effects by binding to specific genomic targets, including *CDH2*, encoding N-cadherin, which emerged as a driver of enhanced migration upon SAMD1 knockout. Furthermore, we discovered the FBXO11-containing E3 ubiquitin ligase complex as an interactor of SAMD1. FBXO11 ubiquitinates SAMD1 within its DNA-binding winged helix domain and inhibits SAMD1 chromatin binding genome-wide. High *FBXO11* expression in PDAC is associated with poor prognosis and increased expression of EMT-related genes, underlining an antagonistic relationship between SAMD1 and FBXO11. In summary, our findings provide new insights into the regulation of EMT-related genes in PDAC, shedding light on the intricate role of SAMD1 and its interplay with FBXO11 in this cancer type.

## Introduction

Pancreatic cancer is a highly lethal form of cancer, accounting for 2.8% of newly diagnosed cancer cases but contributing to 4.7% of cancer-related deaths (Sung et al., 2021). Unlike many other cancer types, the incidence and mortality rates have steadily increased in recent years (Cronin et al., 2022). The most prevalent and severe subtype of pancreatic cancer is pancreatic ductal adenocarcinoma (PDAC), with a 5-year survival rate of only 9% in the US (Cronin et al., 2022).

Due to unspecific and late-occurring symptoms, PDAC is usually diagnosed in advanced stages (Oberstein and Olive, 2013). Treatment options mainly include surgery and chemotherapy; however, most tumors are already deemed inoperable at diagnosis (Kleeff et al., 2016). Epithelial-mesenchymal transition (EMT) is a crucial process in pancreatic cancer and is involved in early metastasis (Nieto et al., 2016). Genetically, PDAC is characterized by diverse mutations, with commonly affected genes including *TP53*, *CDKN2A*, *SMAD4*, and *KRAS* (Samuel and Hudson, 2011). These genetic alterations contribute to the heterogeneity of tumors, which can vary substantially from patient to patient (Samuel and Hudson, 2011). Recent advancements in PDAC research have focused on targeting the tumor microenvironment (TME). The TME in PDAC is characterized by its dense and desmoplastic nature, thereby influencing the druggability and chemoresistance of the tumor (Uzunparmak and Sahin, 2019). However, the identification of novel biomarkers for the early diagnosis of PDAC and the investigation of new druggable proteins are needed.

Sterile alpha motif domain-containing protein 1 (SAMD1) belongs to a novel class of CpG island-binding proteins alongside the histone acetyltransferases KAT6A and KAT6B (Stielow et al., 2021a; Stielow et al., 2021b; Weber et al., 2023). These proteins share a winged-helix (WH) domain that enables direct interaction with unmethylated CpG-rich DNA (Stielow et al., 2021b; Weber et al., 2023). CpG islands (CGIs) are regulatory elements commonly found at promoter regions and play a critical regulatory role. Methylation of CGIs typically results in transcriptional silencing, while unmethylated CGIs are associated with active gene transcription (Deaton and Bird, 2011). CXXC domain-containing proteins and Polycomb-like proteins (PCLs) have also been identified to interact specifically with unmethylated CGIs (Fischer and Liefke, 2023; Li et al., 2017; Thomson et al., 2010; Voo et al., 2000).

In mouse embryonic stem cells, SAMD1 was found to be present at thousands of unmethylated CGIs and to recruit the chromatin regulator L3MBTL3 and the histone demethylase KDM1A to its genomic targets, thereby acting as a transcriptional repressor (Stielow et al., 2021b). Deletion of SAMD1 in mouse embryonic stem cells leads to the dysregulation of multiple cellular pathways, including neuronal, developmental, and immune response pathways (Stielow et al., 2021b). The absence of SAMD1 during mouse embryogenesis primarily impairs brain development and angiogenesis and leads to embryonic lethality (Campbell et al., 2023). SAMD1 function is also linked to muscle adaptation after exercise (Dungan et al., 2022) and autism spectrum disorders (Annear et al., 2022).

In multiple tumor types, *SAMD1* expression is upregulated (Stielow et al., 2021a), and in liver cancer cells, knockout of SAMD1 has been shown to reduce proliferation and clonogenicity (Simon et al., 2022). Moreover, liver cancer patients with high levels of SAMD1 exhibit a more unfavorable transcriptional network (Simon et al., 2022). In the context of other cancer types, the role of SAMD1 remains largely unexplored.

Here, we show that in PDAC, SAMD1 acts as a repressor of EMT-related genes. After deleting SAMD1 in PDAC cell lines, we observed increased migration rates and upregulation of cancer-associated pathways, including the EMT pathway. We identified *CDH2* as a key downstream target of SAMD1 that is important for the migration phenotype. Furthermore, we identified the E3 ubiquitin ligase F-box only protein 11 (FBXO11) as an interactor of SAMD1 in PDAC, which inhibits the chromatin association of SAMD1. Together, our study offers novel insights into the control of EMT-related genes in PDAC, revealing the intricate involvement of SAMD1 and its interplay with FBXO11 in this cancer type.

## Results

### SAMD1 regulates EMT pathways in PDAC

Investigation of public cancer gene expression data from TCGA showed that *SAMD1* is commonly upregulated in cancer (Tang et al., 2017; Weinstein et al., 2013) (**Supplementary Figure 1a**). In some cancer types, high *SAMD1* expression correlates with poor prognoses, such as in liver cancer (LIHC) and kidney renal clear cell carcinoma (KIRC), indicating a more oncogenic role (**Supplementary Figure 1b**). In some other cancer types, such as cervical cancer (CESC) and thymoma (THYM), high *SAMD1* expression correlates with a better prognosis (**Supplementary Figure 1c**), suggesting a more tumor-suppressive role. Interestingly, the distinct relationships to patient survival do not correlate with changes in *SAMD1* gene expression upon tumorigenesis, given that *SAMD1* has increased expression in most cancer tissues compared to normal tissues (**Supplementary Figure 1a**). Thus, it is currently unknown why *SAMD1* has these potentially opposing roles in different cancer types.

A vital cancer type in which high *SAMD1* expression correlates with a better prognosis is pancreatic ductal adenocarcinoma (PDAC) (**Figure 1a**). We hypothesized that gaining a deeper understanding of the potential tumor-suppressive role of SAMD1 in PDAC may allow us to employ this function to limit cancer growth. To acquire insights into the role of SAMD1 in PDAC, we investigated TCGA data using gene set enrichment analysis (GSEA) (Subramanian et al., 2005) and compared PDAC samples with high and low *SAMD1* expression. In the samples with high *SAMD1* expression, we found a strongly decreased expression of genes related to epithelial-mesenchymal transition (EMT) (**Figure 1b**). Similar results were also obtained for thymoma and cervical cancer (**Supplementary Figure 1d**), where high *SAMD1* expression also correlates with a better prognosis (**Supplementary Figure 1b**).

**Figure 1:**
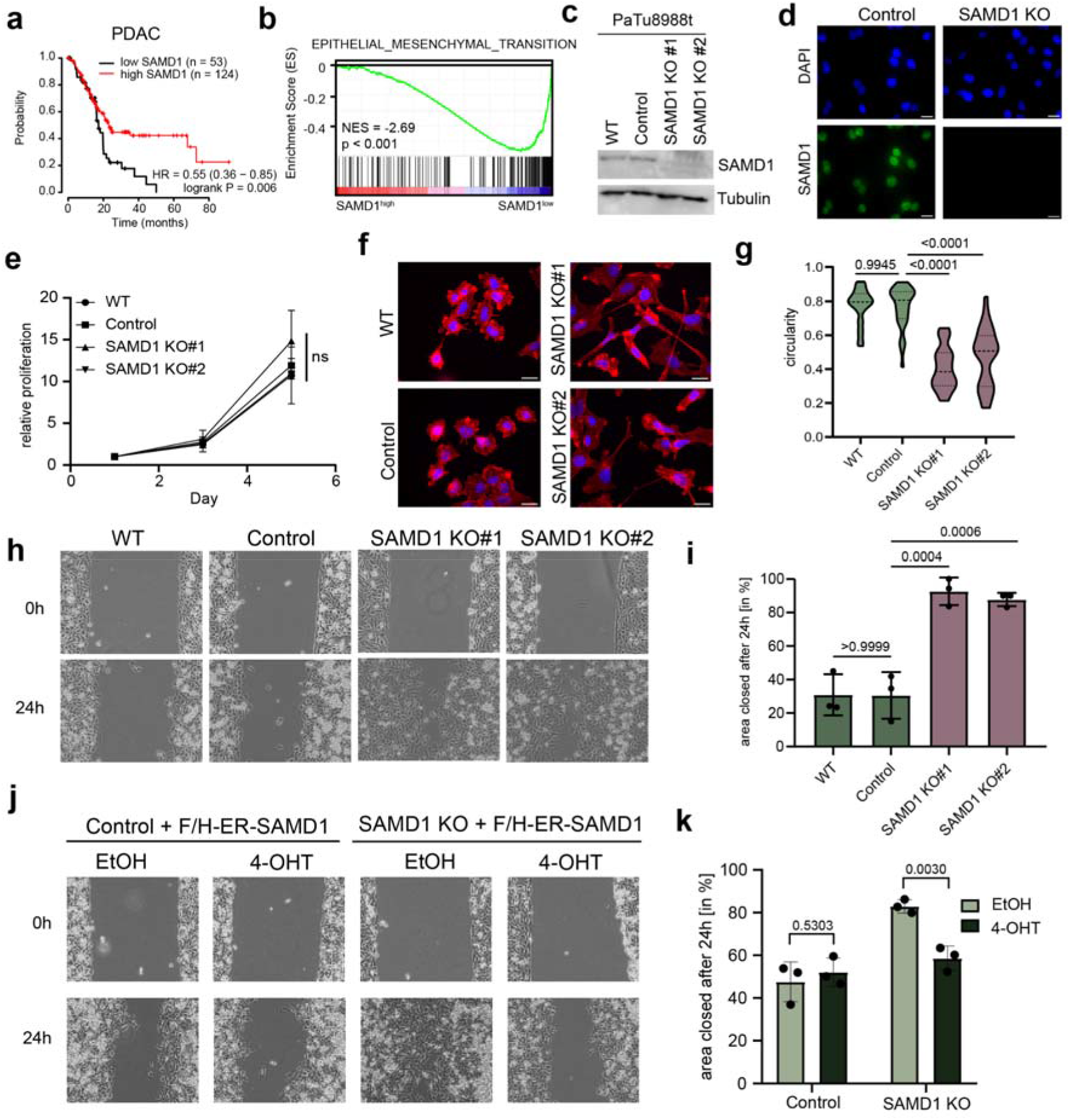
SAMD1 inhibits EMT pathways and cell migration. a) Kaplan Meier survival curves showing the correlation of *SAMD1* expression with patient survival. Blots were visualized via KM-Plotter (Nagy et al., 2021). b) GSEA for epithelial-mesenchymal transition using TCGA data analyzed for high and low *SAMD1* expression. c) Western blot showing PaTu8988t wild-type cells, control cells, and two different SAMD1 knockout clones. d) Immunofluorescence of PaTu8988t wild-type, and SAMD1 knockout cells, Bar=20 µM. e) Proliferation assay of PaTu8988t wild-type, control cells and two different SAMD1 knockout clones, Data represent the mean ± SD of three biological replicates. Significance was analyzed using one-way ANOVA. f) Representative phalloidin staining of PaTu8988t wild-type cells, control cells and two different SAMD1 knockout clones, Bar=20 µM. g) Cell shape of PaTu8988t wild-type cells, control cells, and two different SAMD1 knockout clones. Circularity was determined using ImageJ Fiji. Significance was analyzed using one-way ANOVA. h) Representative picture of one wound healing assay of PaTu8988t wild-type cells, control cells, and two different SAMD1 knockout clones. i) Quantification of the wound healing assay from h). Data represent the mean ± SD of three biological replicates. Significance was analyzed using one-way ANOVA. j) Representative picture of one wound healing assay of PaTu8988t control or SAMD1 KO cells with or without induction of SAMD1 rescue. k) Quantification of the wound healing assay from m). Data represent the mean ± SD of three biological replicates. Significance was analyzed using Student’s t-test.

In contrast, the opposite is the case in cancer types where high *SAMD1* correlates with a worse prognosis, such as kidney cancer. Here, the expression of EMT pathway genes positively correlates with *SAMD1* expression (**Supplementary Figure 1b, d**). Notably, in all investigated cancer types, high *SAMD1* expression correlates with high expression of MYC target genes (**Supplementary Figure 1e**), independent of whether high *SAMD1* expression is favorable or unfavorable, suggesting that this feature is not predictive. Together, these observations lead to the hypothesis that in some cancer types, such as PDAC, SAMD1 may be involved in repressing EMT pathways, thereby inhibiting metastasis, which in turn may contribute to a better outcome. In PDAC, EMT is particularly relevant because it strongly correlates with the occurrence of metastasis, which massively reduces the chance of survival (Wang et al., 2018).

To address the role of SAMD1 in the PDAC cells in more detail, we performed CRISPR/Cas9-mediated knockout approaches in the PDAC cell line PaTu8988t, validated via Western and immunofluorescence (**Figure 1c, d**). We did not observe a change in the proliferation rate upon SAMD1 deletion (**Figure 1e**). Still, investigation of the cells under the microscope showed that SAMD1 knockout led to a more elongated cell shape and more pronounced protrusions (**Figure 1f, g**). This phenotype suggested an increased mobility of the knockout cells. By analyzing the migration rates of PaTu8988t cells by wound healing (**Figure 1h, i**), we confirmed the higher mobility of the SAMD1 KO cells. Furthermore, transwell migration through 8 µm pores demonstrated increased migration rates after SAMD1 deletion in PaTu8988t cells (**Supplementary Figure 2a**), which could be visualized via crystal violet staining (**Supplementary Figure 2b**). We confirmed these results via an unbiased approach by tracking PaTu8988t cells for 24 h using time-lapse analysis (**Supplementary Figure 2c**, **Supplementary Video 1 and 2**).

Enhanced cellular migration, but no chance in proliferation, could also be found in BxPC3 PDAC cancer cells upon SAMD1 deletion (**Supplementary Figure 2d**-g). These results suggest that the influence on the cellular properties by SAMD1 is not restricted to a single cell line but is a more general theme in PDAC. This is further supported by the anticorrelation of *SAMD1* expression and EMT pathways in PDAC patients (**Figure 1b**).

To investigate whether the observed phenotype depends on the nuclear function of SAMD1, we made use of an estrogen-receptor (ER)-SAMD1 fusion protein, whose nuclear localization can be induced by 4-hydroxy-tamoxifen (4-OHT) (**Supplementary Figure 3a, b**). Using this approach, we demonstrated that the migration phenotype in PaTu8998t SAMD1 KO cells can be rescued upon translocation of SAMD1 into the nucleus (**Figure 1j, k**), suggesting that the observed phenotype is linked to the chromatin regulatory role of SAMD1.

### SAMD1 directly represses CDH2, a key regulator of EMT

To address which SAMD1 target genes participate in this phenotype, we performed gene expression analysis via RNA-Seq upon SAMD1 knockout and analyzed the genomic distribution of SAMD1 via ChIP-Seq in PaTu8998t cells. Principal component analysis (PCA) of the RNA-Seq data demonstrated that the knockout led to a substantial shift in the transcriptional landscape (**Supplementary Figure 4a**). The knockout of SAMD1 led to significant deregulation of 854 genes, with 642 upregulated and 212 downregulated genes (cut-off: log2-fold-change > 0.5; p-value < 0.01) (**Figure 2a, Supplementary Figure 4b**). GSEA analysis of the RNA-Seq data demonstrated that the deletion of SAMD1 leads to the dysregulation of multiple cancer-related pathways (**Figure 2b**). Specifically, we observed an upregulation of signaling pathways, including Hedgehog, KRAS, and WNT, and a downregulation of MYC and E2F target genes. Additionally, many transcription factors, including HOXB cluster genes, become dysregulated upon SAMD1 deletion (**Supplementary Figure 4b**). The EMT pathway genes were upregulated in the knockout cells as well (**Figure 2c**), consistent with the observed phenotype and in line with our initial hypothesis that SAMD1 may be involved in regulating this pathway.

**Figure 2:**
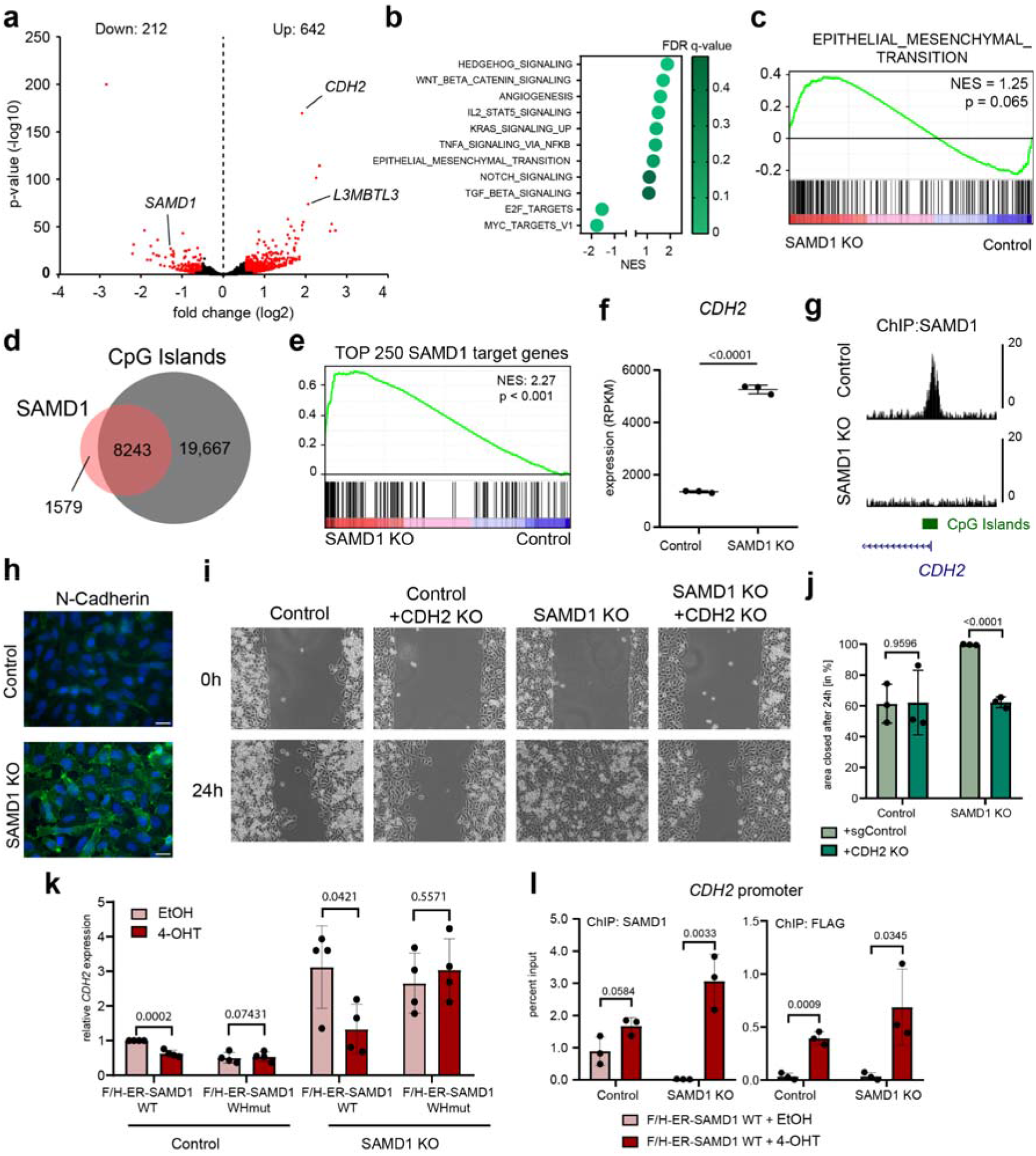
SAMD1 directly regulates EMT pathway genes in PaTu8988t cells. a) Volcano plot of RNA-seq data comparing the results from three replicates of PaTu8988t control cells with three clonally independent SAMD1 KO cells. b) GSEA for several pathways comparing the results from three replicates of PaTu8988t control cells with three clonally independent SAMD1 KO cells. c) GSEA of epithelial-mesenchymal transition from b). d) Venn diagram showing the overlap of SAMD1 peaks with all CpG islands in PaTu8988t cells. e) GSEA of the top 250 SAMD1 targets in PaTu8988t cells, comparing the results from three replicates of control cells with three clonally independent SAMD1 KO cells. f) RNA-Seq results for *CDH2* expression (RPKM) comparing the results from three replicates of PaTu8988t control cells with three clonally independent SAMD1 KO cells. g) Snapshot of the USCS browser showing a SAMD1 peak at the *CDH2* promoter in PaTu8988t control and SAMD1 KO cells. h) Immunofluorescence of N-cadherin in PaTu8988t control and SAMD1 KO cells, Bar=20 µM. i) Representative picture of one wound healing assay of PaTu8988t control, CDH2 KO, SAMD1 KO and CDH2/SAMD1 double KO cells. j) Quantification of the wound healing assay from i). Data represent the mean ± SD of three biological replicates. Significance was analyzed using Student’s t-test. k) RT-qPCR showing *CDH2* expression with or without induction of SAMD1 rescue in PaTu8988t Control and SAMD1 KO cells. WHmut=RK-45/46-AA mutation of *SAMD1*. Data represent the mean ± SD of four biological replicates. Significance was analyzed using Student’s t-test. i) SAMD1 ChIP-qPCR at the *CDH2* promoter with or without induction of SAMD1 rescue in PaTu8988t Control and SAMD1 KO cells. Data represent the mean ± SD of three biological replicates. Significance was analyzed using Student’s t-test.

Genome-wide analysis of SAMD1 chromatin binding showed that SAMD1 mainly binds to CpG island-containing gene promoters (**Figure 2d**), predominantly linked to chromatin and transcriptional regulation (**Supplementary Figure 4c**). The top SAMD1 target genes were significantly upregulated upon SAMD1 deletion, as assessed by GSEA (**Figure 2e**). These results are consistent with our previous findings from mouse ES (Stielow et al., 2021b) and HepG2 cells (Simon et al., 2022) and support that SAMD1 acts as a repressor at CGIs. Consequently, we hypothesized that the increased migratory ability of the knockout cells may be established by the derepression of one or several SAMD1 target genes.

One of the top upregulated EMT genes in the SAMD1 KO cells was *CDH2* (**Figure 2a, f**), whose promoter also showed high SAMD1 enrichment (**Figure 2g**). *CDH2* encodes for N-cadherin, a crucial regulator of cell adhesion and consequently for EMT in PDAC (Derycke and Bracke, 2004; Nakajima et al., 2004). We also confirmed the upregulation of N-cadherin via immunofluorescence (**Figure 2h**). Therefore, *CDH2* could be a key downstream target of SAMD1 in PaTu8988t cells to influence cellular migration. Indeed, the cells possessing a double knockout of CDH2 and SAMD1 had similar characteristics to the wild-type cells regarding their migratory ability (**Figure 2i, j**) and cellular shape (**Supplementary Figure 5a, b**). Additionally, short-term inhibition of N-cadherin-mediated cell adhesion by ADH-1 (Exherin) (Shintani et al., 2008) reduced the elongated phenotype of SAMD1 KO cells, making them more similar to wild-type cells (**Supplementary Figure 5c, d**). These findings support that *CDH2* is a critical downstream factor of SAMD1 that regulates the migration properties of the PaTu8988t cells.

To assess whether SAMD1 directly regulates *CDH2*, we used the ER-SAMD1 fusion protein described above. In SAMD1 KO cells expressing this fusion protein, the expression of *CDH2* was rescued upon 4-OHT treatment (**Figure 2k**). This rescue does not work with a winged-helix domain mutant of SAMD1, indicating that the chromatin binding of SAMD1 is essential for the repression of *CDH2*. Furthermore, ChIP-qPCR confirmed that SAMD1 chromatin binding to the *CDH2* promoter can be rescued with the ER-SAMD1 fusion protein (**Figure 2l**). Similar results were also obtained for the *L3MBTL3* gene, a known SAMD1 target gene that becomes consistently upregulated upon SAMD1 deletion in several distinct cell types (Simon et al., 2022; Stielow et al., 2021b) (**Supplementary Figure 3c, d**). The regulatory effect of SAMD1 on *CDH2* and *L3MBTL3* was also be confirmed in BxPC3 cells (**Supplementary Figure 2h, i**).

Together, these results suggest that SAMD1 is directly involved in repressing *CDH2* in PDAC cells, an essential regulator of EMT (Nakajima et al., 2004).

### The repressive activity of SAMD1 likely involves KDM1A

The molecular details of the repressive function of SAMD1 are currently not fully understood. Previously, we showed that SAMD1 interacts with the KDM1A complex, which demethylates the active H3K4me2 histone mark, and the SAM- and MBT-domain proteins L3MBTL3 and SFMBT1 (Stielow et al., 2021b). In the context of PaTu8988t cells, we confirmed that in the absence of SAMD1, the levels of KDM1A and L3MBTL3 are reduced at the *CDH2* gene, which can be rescued upon induction of the ER-SAMD1 with 4-OHT (**Figure 3a**). Consistently, via ChIP-qPCR, we observed an increase in H3K4me2 and H3K4me3 at the *L3MBTL3* and *CDH2* gene promoters upon SAMD1 deletion (**Figure 3b**). We also validated the interaction between SAMD1 and KDM1A in PaTu8988t cells via endogenous co-immunoprecipitation (**Figure 3c**). These results support that in PaTu8988t cells, KDM1A is likely involved in the repressive function of SAMD1.

**Figure 3:**
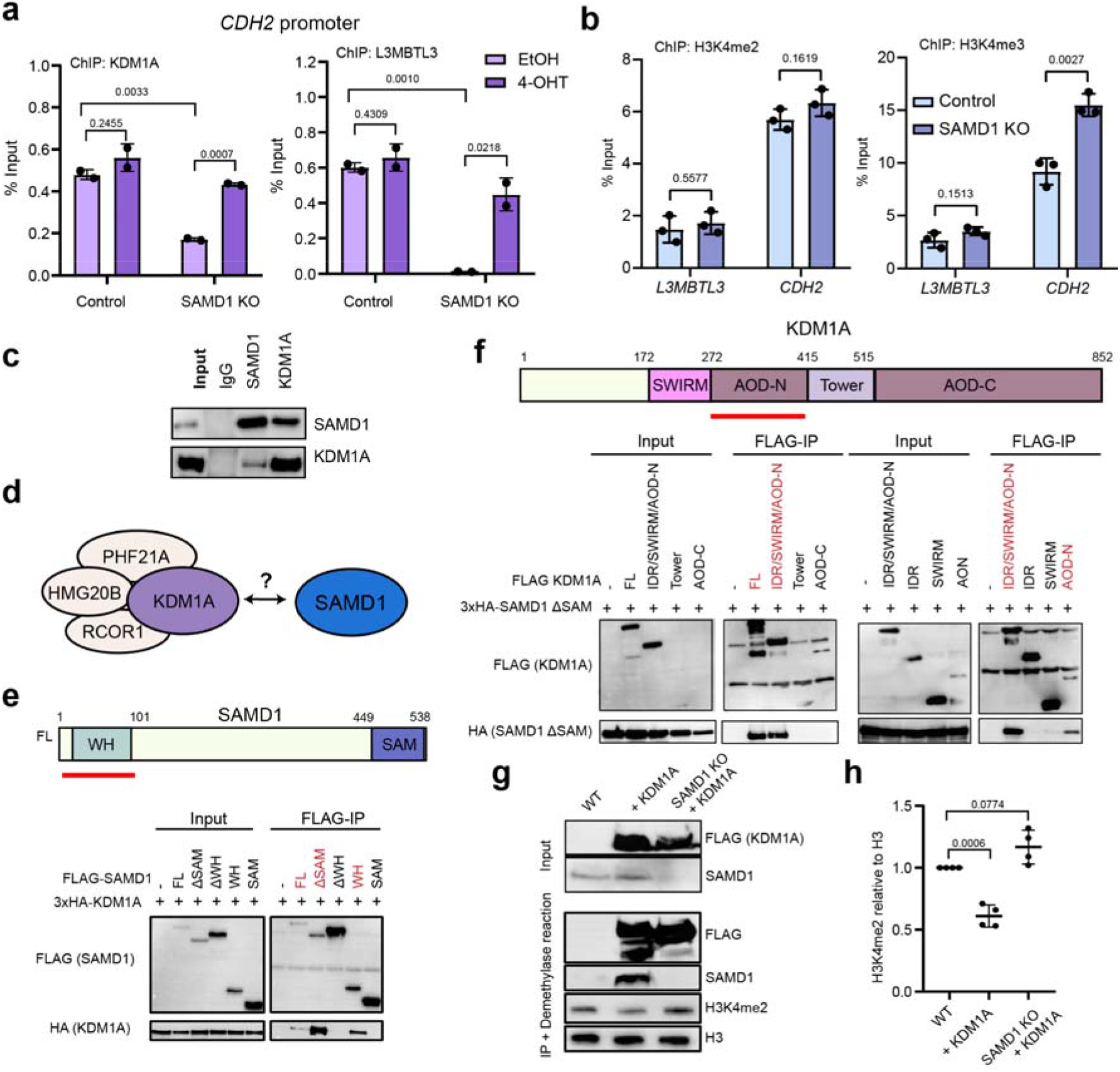
SAMD1 is required for full activity of KDM1A. a) ChIP-qPCR at the *CDH2* promoter with or without induction of SAMD1 rescue in PaTu8988t Control and SAMD1 KO cells using KDM1A and L3MBTL3 antibodies. Data represent the mean ± SD of two biological replicates. Significance was analyzed using Student’s t-test. b) ChIP-qPCR of *CDH2* and *L3MBTL3* promoter in PaTu8988t Control and SAMD1 KO cells using H3K4me2 and H3K4me3 antibodies. Data represent the mean ± SD of three biological replicates. Significance was analyzed using Student’s t-test. c) Western blot of an endogenous Co-IP between SAMD1 and KDM1A in PaTu8988t cells. d) Model of the interaction between SAMD1 and the KDM1A complex. e) Structure of SAMD1; co-immunoprecipitation in HEK293 cells showing the interaction between different SAMD1 deletion mutants and KDM1A. Regions identified to interact with KDM1A are marked red. f) Structure of KDM1A; co-immunoprecipitation in HEK293 cells showing the interaction between different KDM1A deletion mutants and SAMD1. Regions identified to interact with SAMD1 are marked red. g) Representative western blot of KDM1A IP in HEK293 cells, followed by histone demethylase assay. h) Quantification of four biological replicates of g). Significance was analyzed using one-way ANOVA.

To gain further insight into the interplay of SAMD1 with the KDM1A complex (**Figure 3d**), we performed mapping experiments that went beyond our previous experiments (Stielow et al., 2021b). First, we confirmed that SAMD1 preferentially interacts with KDM1A via its winged-helix domain and that deleting the SAM domain, which is essential for the interaction with L3MBTL3 (Stielow et al., 2021b), increases the interaction with KDM1A (**Figure 3e**). This phenomenon is also observable with other members of the KDM1A complex (**Supplementary Figure 6a**). Reverse mapping experiments suggested that SAMD1 preferentially interacts with the N-terminal part of the catalytic amino oxidase domain (AOD) of KDM1A (**Figure 3f**). This result is further supported by the observation that the KDM1A inhibitor ORY-1001 (Iadademstat), which covalently binds to the FAD cofactor within KDM1A (Maes et al., 2018), interferes with the interaction of KDM1A with SAMD1 (**Supplementary Figure 6b**). This effect was not observed for other KDM1A complex members, such as RCOR1 and PHF21A (**Supplementary Figure 6c, d**). This finding raises the possibility that SAMD1 may bind near the catalytic cleft of KDM1A, making the interaction sensitive to the inhibitor treatment.

Based on these results, we speculated that SAMD1 could potentially influence the demethylase activity of KDM1A, similar to other factors associated with the KDM1A complex (Shi et al., 2005). To address this question, we immunoprecipitated KDM1A in either wild-type or SAMD1 KO HEK293 cells and used the obtained precipitate for demethylase assays, adding calf histones as a substrate. We found that the absence of SAMD1 reduced the ability of KDM1A to remove H3K4me2 efficiently (**Figure 3g, h**). This result suggests that SAMD1 modulates the function of KDM1A, possibly not just by influencing its recruitment to chromatin but also by influencing the catalytic activity of KDM1A. Both processes together may contribute to the repressive role of SAMD1. However, the molecular details of how SAMD1 affects the enzymatic activity of KDM1A require further research. In addition, we cannot exclude the possibility that other mechanisms, such as the recruitment of L3MBTL3, are also crucial for the repressive function of SAMD1.

### SAMD1 interacts with the FBXO11 E3 ubiquitin ligase complex

Upon investigating the cellular localization of SAMD1 in PaTu8988t cells and further PDAC cell lines, we observed that compared to other human cell lines, SAMD1 is less present in the chromatin fraction in PDAC cells (**Figure 4a**). This finding raises the possibility that a certain molecular mechanism regulates the chromatin binding of SAMD1. In the context of pancreatic cancer cells, such a mechanism may be essential to overcome the tumor-suppressive function of SAMD1. To date, no process has been described that regulates the chromatin association of SAMD1.

**Figure 4:**
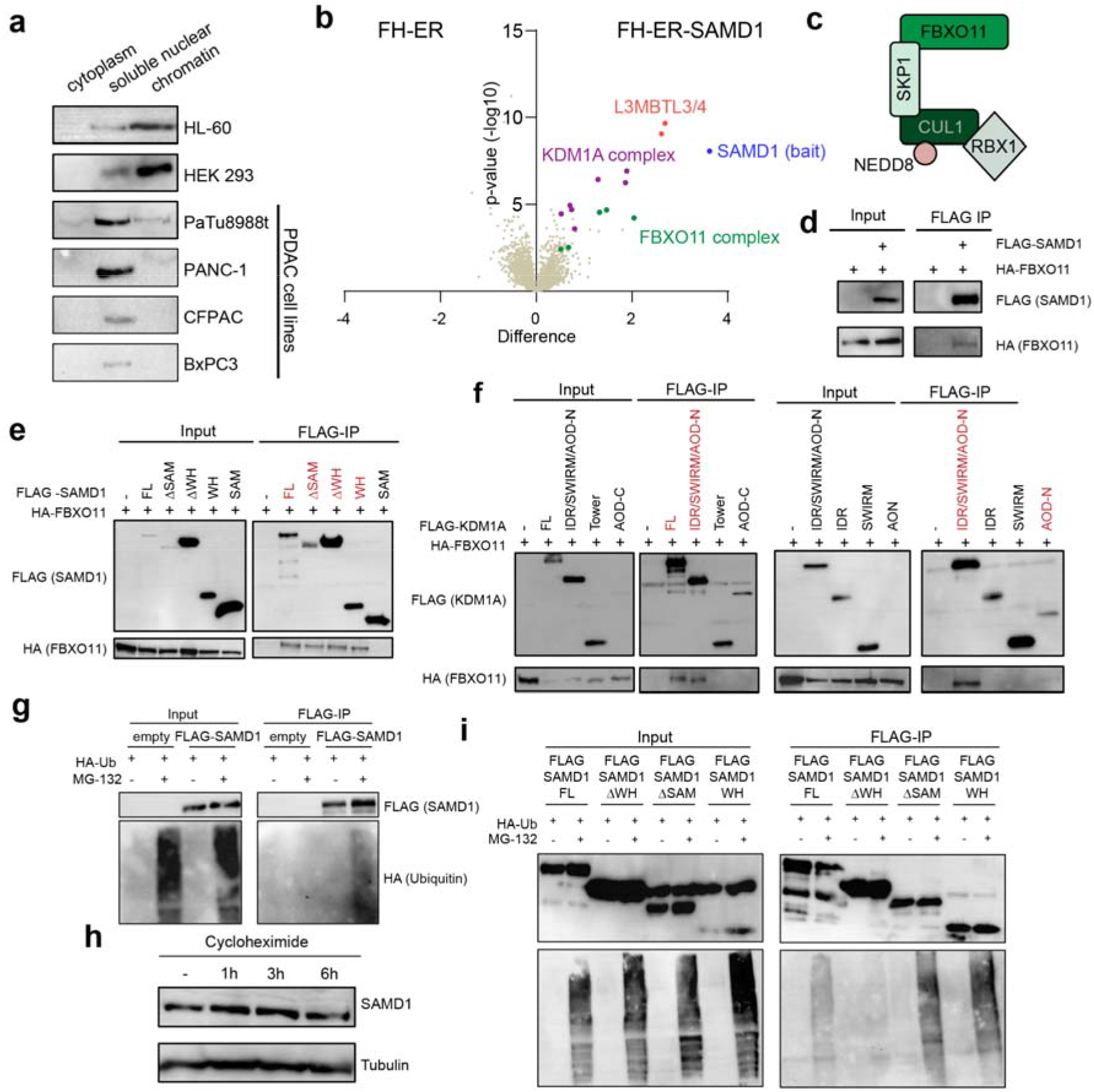
SAMD1 interacts with FBXO11 in PaTu8988t cells and is ubiquitinated. a) Fractionation of different cell lines followed by SAMD1 western blotting. b) Volcano plot of proteins identified by mass spectrometry after IP of FH-ER and FH-ER-SAMD1. c) Model of the FBXO11 E3-ubiquitin ligase complex. d) Co-immunoprecipitation in HEK293 cells showing the interaction between SAMD1 and FBXO11. e) Co-immunoprecipitation in HEK293 cells showing the interaction between different SAMD1 deletion mutants and FBXO11. f) Co-immunoprecipitation in HEK293 cells showing the interaction between different KDM1A deletion mutants and FBXO11. g) Ubiquitination assay after empty vector or SAMD1 transfection in HEK293 cells. h) Cycloheximide chase analysis of SAMD1 in PaTu8988t cells. i) Ubiquitination assay in HEK293 cells using different SAMD1 deletion constructs.

To address whether SAMD1 may interact with additional proteins that could be involved in such a regulatory process, we performed unbiased IP-MS experiments in PaTu8988t cells. For this, we used cells expressing the ER-SAMD1 fusion protein (**Supplementary Figure 3a, b**). After inducing the nuclear localization of the protein via 4-OHT, we collected the cells and immunoprecipitated the SAMD1 protein. The cobound proteins were analyzed by LC-MS (**Figure 4b**). This experiment confirmed that SAMD1 interacts with L3MBTL3 and the KDM1A histone demethylase complex. We also identified L3MBTL4, consistent with our finding that the SAM domain of L3MBTL4 can interact with the SAM domain of SAMD1, similar to L3MBTL3 (Stielow et al., 2021b). In addition to these expected interactions, we identified members of the FBXO11 complex as putative novel interaction partners of SAMD1. The FBXO11 complex consists of FBXO11 itself, RBX1, Cullin 1, and SKP1, all of which are enriched in the SAMD1 IP (**Figure 4b, c**). Additionally, we found enrichment of NEDD8, which is typically covalently associated with Cullin 1 and is required for the F-box protein-associated E3 ubiquitin ligase complexes to be active (Zheng et al., 2002).

Via co-immunoprecipitation experiments in HEK293 cells, we validated that SAMD1 can interact with FBXO11 (**Figure 4d**). Additional mapping experiments suggested that several regions of SAMD1 are relevant for this interaction (**Figure 4e**). Only the SAM domain appears dispensable for the interaction with FBXO11 (**Figure 4e**). Interestingly, we found that FBXO11 can also be co-immunoprecipitated with KDM1A (**Figure 4f**). This interaction is facilitated by the N-terminal part of the AOD domain of KDM1A, which is the same region as for the interaction with SAMD1 (**Figure 4f, 3f**). Given that the interaction between KDM1A and FBXO11 is also sensitive to the KDM1A inhibitor ORY-1001 (**Supplementary Figure 6e**), similar to the SAMD1/KDM1A interaction (**Supplementary Figure 6b**), it suggests that KDM1A interacts with SAMD1 and FBXO11 simultaneously.

FBXO11 is an E3 ubiquitin ligase that regulates the ubiquitination of various target proteins, including BCL6, CDT2, and BAHD1 (Duan et al., 2011; Rossi et al., 2013; Xu et al., 2020). We hypothesized that FBXO11 may facilitate the ubiquitination of SAMD1. To investigate whether SAMD1 is ubiquitinated, we co-immunoprecipitated FLAG-tagged SAMD1 in HEK293 cells expressing HA-tagged ubiquitin. We observed that in the presence of the proteasome inhibitor MG-132, the immunoprecipitated FLAG-SAMD1 but not the control precipitate showed an HA-signal in Western blotting experiments (**Figure 4g**), demonstrating that SAMD1 becomes ubiquitinated when the proteasome is inhibited. Surprisingly, however, cycloheximide chase experiments suggested that SAMD1 is highly stable, with no obvious turn-over within 6 hours (**Figure 4h**). To assess which region of SAMD1 is mostly ubiquitinated, we used SAMD1 deletion mutants. We found that deleting the WH domain decreased the ubiquitination level, while deletion of the SAM domain, or using the WH domain alone led to an increased ubiquitination level (**Figure 4i**). This result suggests that the WH domain is the primary ubiquitination site of SAMD1.

### Ubiquitination by FBXO11 possibly directly influences the DNA binding of SAMD1

To investigate the role of FBXO11 in the ubiquitination of SAMD1, we established HEK293 and PaTu8988t cells with FBXO11 knockout (**Figure 5a**). Consistent with the idea that FBXO11 ubiquitinates SAMD1, we found a reduced level of ubiquitination of the SAMD1 WH domain in the FBXO11 knockout HEK293 cells (**Figure 5b, c**).

**Figure 5:**
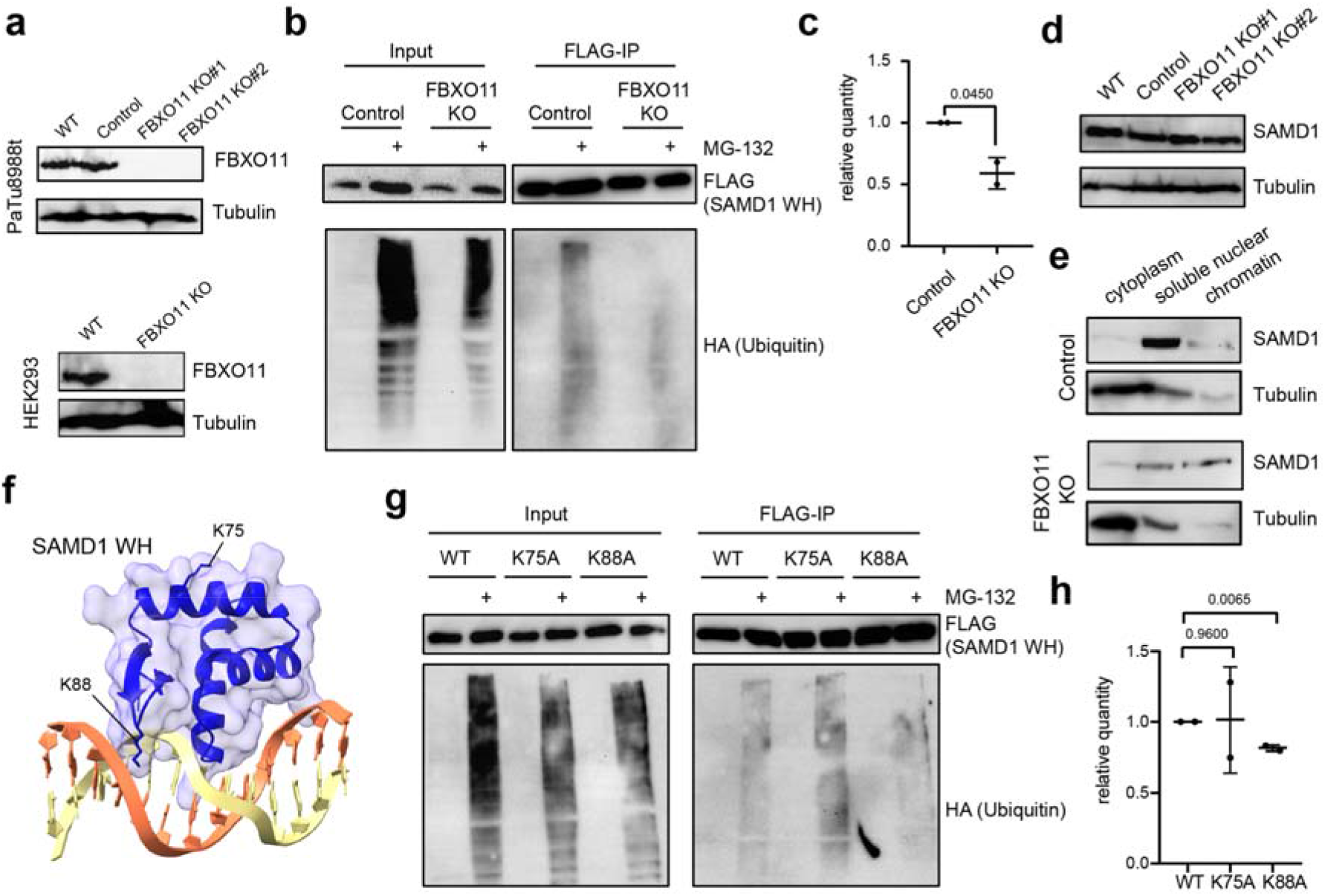
FBXO11 affects SAMD1 ubiquitination and chromatin association. a) Western blot showing PaTu8988t wild-type cells, control cells, and two different FBXO11 knockout clones; western blot showing HEK293 cells with FBXO11 KO. b) Ubiquitination assay of SAMD1 winged helix (WH) domain in HEK293 control and FBXO11 KO cells. c) Quantification of b) using two biological replicates. d) Western blot showing SAMD1 expression in PaTu8988t wild-type cells, control cells, and two different FBXO11 knockout clones. e) Fractionation of PaTu8988t control and FBXO11 KO cells, followed by SAMD1 western blotting. f) Structure of the SAMD1 WH domain (PDB: 6LUI)(Stielow et al., 2021b), with K75 and K88 indicated. g) Ubiquitination assay of wild-type and mutated SAMD1 winged-helix domains in HEK293 cells. h) Quantification of g) using two biological replicates.

Notably, in the FBXO11 knockout PaTu8988t cells, we did not observe an altered protein level of SAMD1 (**Figure 5d**), suggesting that ubiquitination of SAMD1 by FBXO11 has no relevant influence on the turn-over of SAMD1. However, fractionation experiments showed that the chromatin association of SAMD1 was substantially augmented in the FBXO11 KO cells (**Figure 5e**), supporting that the FBXO11 is involved in modulating the chromatin association of SAMD1.

We speculated that the inhibitory effect of FBXO11 on SAMD1 chromatin binding is due to direct interference with SAMD1’s DNA binding ability. Based on the PhosphoSitePlus database (Hornbeck et al., 2015), the amino acids K75 and K88, which lie in the DNA-binding WH domain, can be ubiquitinated. K88 is directly involved in DNA binding via its ability to interact with the minor groove of the DNA (Stielow et al., 2021b), while K75 is not associated with DNA (**Figure 5f**). We mutated these amino acids into alanines to address whether they are relevant for SAMD1 ubiquitination. Indeed, we found an approximately 30% percent reduction in ubiquitination levels with the K88A mutant, suggesting that this site is important for this process (**Figure 5g, h**).

Ubiquitination of K88 would lead to sterical hindrance and would prevent the formation of H-bridges of K88 with the DNA. Thus, it is likely that this ubiquitination would at least reduce the DNA binding capacity of the WH domain. However, previous experiments showed that K88 is less critical for DNA binding than R44 and K45, which bind to the major groove of the DNA (Stielow et al., 2021b). Also, the homologous WH domain of KAT6A shows only subtle association with the minor groove (Weber et al., 2023), indicating that this interaction is not absolutely required for the DNA binding. Thus, we cannot exclude the possibility that ubiquitination of K88 still allows binding of SAMD1 to the DNA, albeit likely less efficiently.

### FBXO11 deletion influences SAMD1 chromatin binding genome-wide

To address the role of FBXO11 in SAMD1 chromatin binding in further detail, we performed ChIP-qPCR using our SAMD1 antibody. Consistent with the fractionation experiment (**Figure 5e**), we observed an enhanced chromatin binding of SAMD1 at the *CDH2* promoter in the FBXO11 KO PaTu8988t cells (**Figure 6a**). To investigate this effect at the genome-wide level, we performed ChIP-Seq of SAMD1 in wild-type and FBXO11 KO cells. We found an increased SAMD1 binding at many locations in FBXO11 knockout cells (**Figure 6b, Supplementary Figure 7a**), supporting that FBXO11 is involved in inhibiting the chromatin binding of SAMD1 at a global level.

**Figure 6:**
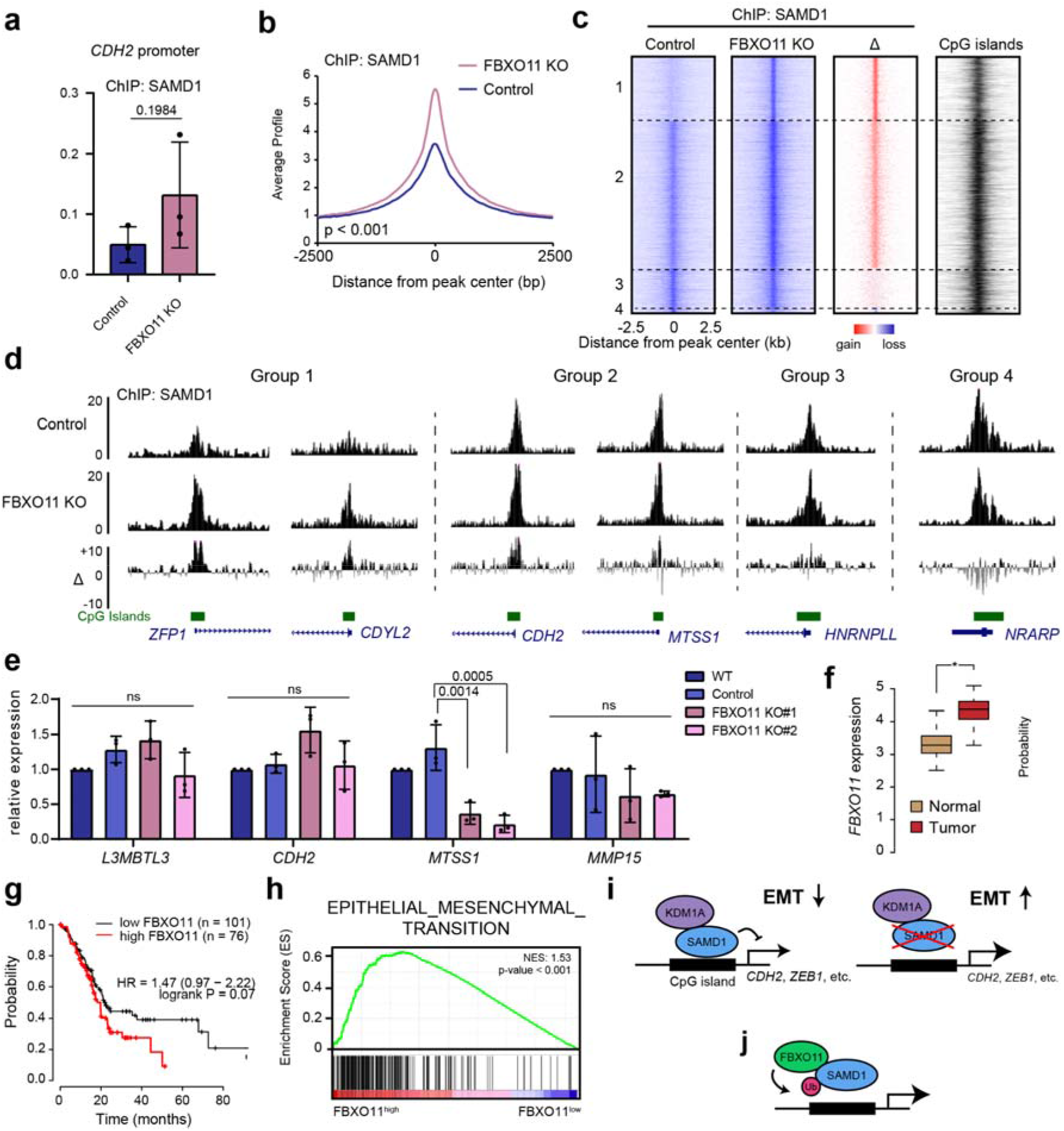
FBXO11 counteracts SAMD1. a) ChIP-qPCR at the CDH2 promoter in PaTu8988t Control and FBXO11 KO cells, using a SAMD1 antibody. Data represent the mean ± SD of three biological replicates. b) Profile of SAMD1 at SAMD1 peaks, in wild-type and FBXO11 KO cells. Significance was evaluated via a Kolmogorov-Smirnov test. See also **Supplementary Figure 7a**. c) Heatmaps showing all SAMD1 peaks in PaTu8988t control and FBXO11 KO cells. Peaks were grouped according to the gain or loss after FBXO11 KO (Group 1, n = 3028; Group 2, n = 6895; Group 3, n = 1883; Group 4, n = 229). Δ indicates the difference between control and FBXO11 KO. Peaks overlap with CpG islands. See also **Supplementary Figure 7b**. d) Snapshots of the USCS browser showing examples of the four groups observed in c). e) RT-qPCR analysis of SAMD1 target genes with enhanced SAMD1 chromatin binding upon FBXO11 KO. Data represent the mean ± SD of three biological replicates. Significance was analyzed using one-way ANOVA. f) Expression of FBXO11 in PDAC versus normal tissues. Data from TCGA (Weinstein et al., 2013) and visualized via GePIA (Tang et al., 2017). g) Kaplan Meier survival curve showing the correlation of *FBXO11* expression with patient survival. Plot was visualized via KM-Plotter (Nagy et al., 2021). h) GSEA for epithelial-mesenchymal transition using TCGA data analyzed for high and low FBXO11 expression. i) Model of the role of SAMD1 in PDAC. SAMD1 represses EMT-related genes, thereby suppressing migration. In SAMD1 KO cells, the EMT genes become upregulated leading to an enhanced EMT-related phenotype. j) FBXO11 is involved in counteracting SAMD1 by ubiquitination, which inhibits SAMD1 chromatin association.

The impact of FBXO11 deletion was particularly evident at locations where SAMD1 chromatin binding was low under wild-type conditions (**Figure 6c, d,** groups 1 and 2, **Supplementary Figure 7b**). In contrast, in places where SAMD1 was already strongly present in the wild-type cells, the FBXO11 knockout had only minor effects (**Figure 6c, d,** group 3, **Supplementary Figure 7b**). Only at a very small fraction, with very high SAMD1 levels, does SAMD1 occupancy become weaker (**Figure 6c, d,** group 4, **Supplementary Figure 7b**). Closer inspection of the distinct groups showed that the locations more susceptible to FBXO11 deletion have smaller CpG islands (**Supplementary Figure 7c**) and are transcriptionally more active, signified by higher H3K4me3 and RNA Polymerase II levels (**Supplementary Figure 7d**-e). This observation raises the possibility that FBXO11 more strongly regulates SAMD1 chromatin binding at locations with fewer CpG binding motifs and higher transcriptional activity. On the other hand, no substantial differences regarding the gene expression levels of the associated genes could be recognized (**Supplementary Figure 7f**).

To investigate whether the enhanced binding of SAMD1 contributes to changes in gene expression, we analyzed several SAMD1 target genes via RT-qPCR. We found a significant reduction in *MTSS1* gene expression but no changes in other investigated genes (**Figure 6e**). This observation suggests that the increased chromatin binding of SAMD1 upon FBXO11 deletion has an impact on the gene expression of specific genes, consistent with the overall relatively low number of dysregulated genes upon SAMD1 deletion (**Figure 2a**).

Our biochemical and genome-wide data support the hypothesis that FBXO11 ubiquitinates SAMD1, which impairs its chromatin association and thereby may inhibit the gene repressive function of SAMD1. This mechanism could be important during cancer progression to overcome the tumor-suppressive role of SAMD1. This idea is further supported by the fact that *FBXO11* is commonly upregulated in PDAC (**Figure 6f**), and its high expression correlates with worse patient prognosis (**Figure 6g**). Furthermore, a high *FBXO11* expression level was linked to increased expression of EMT pathway genes (**Figure 6h**), which is opposite to what we observed before for *SAMD1 (***Figure 1b**). Together, these data support that in PDAC cells, the tumor-suppressive function of SAMD1 is counteracted by FBXO11.

## Discussion

Pancreatic ductal adenocarcinoma (PDAC) is a highly lethal form of cancer (Kleeff et al., 2016). Local invasion and metastasis, driven by uncontrolled epithelial-mesenchymal transition (EMT), are the main reasons for the aggressive nature of PDAC (Wang et al., 2018). Unfortunately, current strategies for inhibiting EMT in PDAC are not sufficiently effective. In this work, we identified the chromatin regulator SAMD1 as an important suppressor of EMT-related pathways in PDAC.

Analysis of patient data revealed that *SAMD1* is frequently dysregulated in cancer, and its expression often correlates with favorable or unfavorable prognoses. In the context of PDAC samples, *SAMD1* expression is commonly upregulated, (**Supplementary Figure 1a**), but Kaplan Meier survival curves demonstrated that high SAMD1 expression is associated with a better outcome (**Figure 1a**), suggesting a tumor-suppressive role for SAMD1. Biological assays utilizing SAMD1 KO cells were conducted to investigate this further, revealing increased migration rates and upregulation of EMT-related pathways upon SAMD1 loss (**Figure 1h, i, Figure 6i**), further supporting a potential tumor-suppressive function of SAMD1.

During this work, we noticed a decreased chromatin binding of SAMD1 in PDAC cell lines compared to other tumor cell lines (**Figure 4a**). This finding led us to speculate that the correlation with survival may not be solely determined by *SAMD1* expression levels but rather by the chromatin-binding level of SAMD1 itself. Via IP-MS experiments we revealed the FBXO11-containing E3-ubiquitin ligase complex as a novel interactor of SAMD1 (**Figure 4b**), which acts as an inhibitor of SAMD1 chromating binding (**Figure 6j**). FBXO11 plays a versatile role in cancer, acting both as an oncogene and as a tumor-suppressor. It targets oncogenic proteins, such as BCL-6 or the Snail family of transcription factors, for degradation, thereby exhibiting a tumor-suppressive function in various cancer types, such as diffuse large B-cell lymphomas and lung cancer (Duan et al., 2011; Jin et al., 2015). In lung cancer cell lines, the FBXO11-containing complex was found to neddylate p53, thereby inhibiting its transcriptional activity (Abida et al., 2007). In contrast, in pancreatic ductal adenocarcinoma, silencing FBXO11 suppresses tumor development (Xue et al., 2022), which may involve the ubiquitination of p53, supporting an oncogenic role in this cancer type. As a result, high *FBXO11* expression is associated with a worse prognosis in PDAC (**Figure 6d**) (Xue et al., 2022). Our finding that FBXO11 inhibits SAMD1 chromatin association represents a novel regulatory mechanism of FBXO11 in cancer (**Figure 6j**). The FBXO11-SAMD1 axis could potentially be relevant for more cancer types beyond PDAC.

In our IP-MS experiments, we successfully identified another key interactor of SAMD1 – the KDM1A histone demethylase complex (**Figure 4b**). KDM1A, also referred to as LSD1, has previously been characterized as an interactor of SAMD1 (Stielow et al., 2021b). In mouse embryonic stem cells, the deletion of SAMD1 results in a reduction of KDM1A-binding on specific promoters (Stielow et al., 2021b), implying that SAMD1 plays a role in the recruitment of KDM1A to chromatin. We confirmed that KDM1A is also an interactor of SAMD1 in PDAC (**Figure 3c**) and that deletion of SAMD1 leads to reduced recruitment of KDM1A to chromatin (**Figure 3a**) and altered H3K4 methylation levels (**Figure 3b**). Besides the involvement of SAMD1 in the recruitment of KDM1A, our work supports the idea that the presence of SAMD1 in the KDM1A complex influences the catalytic activity of KDM1A (**Figure 3g, h, Figure 6i**). One could speculate that the association of SAMD1 with KDM1A affects the conformation of the KDM1A complex, which in turn may allow a more efficient demethylation reaction. Another possibility is that the SAMD1, when bound to the KDM1A complex, enhances the association of the KDM1A complex with its nucleosomal substrate, which increases the efficiency of demethylation. It is also possible that the association of KDM1A with the FBXO11 complex contributes to the altered demethylase activity of KDM1A. Thus, more research will be required to clarify the potentially sophisticated interplay of KDM1A, SAMD1, and FBXO11.

*CDH2* is one of the top SAMD1 target genes and shares occupancy with KDM1A (**Figure 2**, **Figure 3a**). Our investigations have demonstrated that *CDH2*, the gene encoding N-cadherin, serves as the main driver of enhanced migration after SAMD1 KO (**Figure 2j, k**). During EMT progression, cadherins play a central role. The downregulation of epithelial cadherin (E-cadherin), which is critical for adherens junction formation, is accompanied by an elevation in neural cadherin (N-cadherin) expression. This shift contributes to heightened cell mobility and a more mesenchymal phenotype (Loh et al., 2019). While N-cadherin is only detectable in nearly 50% of all PDAC patient samples, its presence in metastatic lesions significantly correlates with augmented neural invasion and a higher histological grade (Nakajima et al., 2004). Remarkably, tumors exhibiting high N-cadherin levels also show elevated *TGFB* expression, which resonates with our observations indicating the upregulation of TGFB-signaling consequent to SAMD1 knockout (**Figure 2b**). Besides *CDH2*, many other EMT-related genes, including *ZEB1 (Zhang et al., 2015), BMP2 (Gordon et al., 2009),* and Netrin-1 (*NTN1*) (Lengrand et al., 2023), are targeted and repressed by SAMD1 (**Figure 6i**).

Based on our RNA-Seq experiment performed here (**Figure 2a**) and previously (Simon et al., 2022; Stielow et al., 2021b), most SAMD1 target genes are only subtly influenced by SAMD1 deletion, leading to less than 10-fold upregulation. However, given that SAMD1 targets thousands of genes (**Figure 2d**), it is likely that SAMD1 has a substantial influence on the transcriptional landscape in the cells. This hypothesis is supported by the intense dysregulation of cellular pathways during differentiation processes (Stielow et al., 2021b) and by the severe phenotype of the SAMD1 knockout mice (Campbell et al., 2023). It is also notable that the impact of SAMD1 on gene transcription is rather cell-type specific. Except for the *L3MBTL3* gene, which appears to be commonly dysregulated upon SAMD1 deletion (**Figure 2a**) (Simon et al., 2022; Stielow et al., 2021b), possibly due to a feedback mechanism, most other genes are affected by SAMD1 in a cell type-specific manner (Simon et al., 2022). Thus, the role of SAMD1 is likely highly context-dependent, and more work will be necessary to fully understand the specific functions of SAMD1 in the various physiological and pathophysiological contexts.

In PDAC, our work identified a tumor-suppressive role of SAMD1 by inhibiting EMT-related genes. By exploiting this functionality, such as enhancing SAMD1 chromatin binding by disrupting its interaction with FBXO11 (**Figure 6j**), it may be possible to impede EMT during PDAC progression. Thus, our findings serve as a foundation for future investigations into the therapeutic possibilities of targeting SAMD1 in PDAC and other diseases.

## Data availability

ChIP-Seq and RNA-Seq data were uploaded to the Gene Expression Omnibus (GEO) database, with the accession numbers GSEXXXXXX and GSEXXXXXX.The mass spectrometry proteomics data have been deposited to the ProteomeXchange Consortium via the PRIDE (Perez-Riverol et al., 2022) partner repository with the dataset identifier PXDXXXXXX.

## Funding

This project was supported by the German Research Foundation (DFG, 109546710, 416910386, 516068166), the German José Carreras Leukemia Foundation (DJCLS 06 R/2022) and the Fritz Thyssen Foundation (10.20.1.005MN) to R.L.

## Author Contributions

C.S. performed most biochemical and biological experiments, I.D.B performed most ubiquitination experiments and created FBXO11 KO cells, B.S. performed most ChIP experiments and prepared ChIP-seq samples, A.M.S. performed transwell migration assays, L.M.J. and M.G. performed mapping experiments between KDM1A and FBXO11, M.B. helped with analysis of time-lapse analysis experiments, I.F. performed mass-spectrometry analysis, I.R. created plasmids for most experiments, M.G, L.M.W. and S.F. contributed experiments and/or material, A.N. performed next-generation sequencing, R.L. performed bioinformatic analyses, T.S., M.H. and R.L supervised the work. C.S. and R.L. conceived the study. R.L. and C.S. wrote the manuscript with input from all authors.

## Acknowledgments

We acknowledge the Protein Analytics Unit at the Biomedical Center, Ludwig-Maximilians University Munich, for providing services and assistance with data analysis. We thank Matthias Lauth and Uta-Maria Bauer for providing PDAC cell lines and for discussions.

## Material and Methods

### Cell culture

Patu8988t cells were cultured in DMEM, high glucose, GlutaMAX™ Supplement (Thermo Fisher Scientific; 61965026) supplemented with 5% fetal bovine serum (Thermo Fisher Scientific; 10270106). PANC-1 and CFPAC cells were cultured in DMEM, high glucose, GlutaMAX™ Supplement supplemented with 10% fetal bovine serum respectively. BxPC3 cells were kept in RPMI 1640 Medium, GlutaMAX™ Supplement (Thermo Fisher Scientific; 61870036) supplemented with 10% fetal bovine serum whereas HL-60 cells obtained 15% fetal bovine serum. HEK 293 cells were grown in DMEM/F-12, GlutaMAX™ Supplement supplemented with 10% fetal bovine serum. HepG2 cells were cultured in MEM, GlutaMAX™ (Thermo Fisher Scientific; 41090036) supplemented with 10% fetal bovine serum and 1xnonessential amino acids (Thermo Fisher Scientific; 11140050). All cell lines were cultured with 1% penicillin-streptomycin (Thermo Fisher Scientific; 15140122).

### Antibodies

All antibodies used are described in the methods subsections and in **Supplementary Table S1**.

### Stable cell line generation

A SAMD1 knockout was conducted using the Lenti-CRISPR V2 plasmid containing either an unspecific control or guide RNAs targeting *SAMD1* (sg1: AGCGCATCTGCCGGATGGTG; sg2: GAGCATCTCGTACCGCAACG), *CDH2* (sg1: GCCTGAAGCCAACCTTAACTG; sg2: GAGACAATTCAGTAAGCACAG; sg3: GAACTTGCCAGAAAACTCCAG), *FBXO11* (sg1: GAGCCTCTTGTACCCCACCA; sg2: GTGTCCCACAAAGAACAGTA; sg3: GTTTTCTGTAGTTGAAGTTG).

Cells were transfected using Polyethylenimine, Linear, MW 25000, Transfection Grade (Polysciences; 23966) and Opti-MEM™ (Thermo Fisher Scientific; 31985062). Selection for single clones was performed using 2 µg/µl puromycin (Merck; 58-58-2) for PaTu8988t cells and 0.3 µg/µl for BxPC3 cells. The knockout was confirmed by western blot or immunofluorescence.

To rescue SAMD1, PaTu8988t cells were transfected with SAMD1 constructs containing a FLAG-HA-ER tag. Positive clones were selected using 10 µg/ml blasticidin. After selection, the concentration was reduced to 5 µg/ml blasticidin. Nuclear translocation of FLAG-ER-SAMD1 was induced by adding 200 nM 4-OHT (Merck; 68392-35-8) for 24 h.

### Nuclear Extract Preparation

To obtain the nuclear extract, the cytoplasmic fraction was removed by incubating harvested cells for 10 min at 4 °C in low salt buffer (10 mM HEPES/KOH (pH=7.9), 10 mM KCl, 1.5 mM MgCl2, 1xPIC (cOmplete™, Protease Inhibitor Cocktail (Roche; 04693116001)), 0.5 mM PMSF). After centrifugation, the remaining pellet was dissolved in high salt buffer (20 mM HEPES/KOH pH=7.9, 420 mM NaCl, 1.5 mM MgCl2, 0.2 mM EDTA, 20% glycerol, 1x PIC, 0.5 mM PMSF) and incubated for 20 min at 4 °C while shaking. Subsequently, the lysates were centrifuged, and the supernatant containing the nuclear fraction was further analyzed by western blotting.

### Subcellular Fractionation

A subcellular protein fractionation kit for cultured cells (Thermo Fisher Scientific; 78840) was used for fractionation experiments according to the manufacturer’s instructions. A 10 cm dish format was applied, which corresponded to a packed cell volume of 20 μl per well.

### Western Blot

Western blots were conducted using the Trans-Blot® Turbo™ Transfer System (BioRad; 1704150). The following antibodies were used: anti-tubulin (Merck; MAB3408), anti-SAMD1 antibody (Bethyl; A303-578A), anti-FBXO11 (Novus Biologicals; NB100-59826), anti-KDM1A (Abcam; AB17721), anti-HA (Merck; 11867423), and anti-FLAG (Merck; F3165).

### Immunofluorescence Staining

For immunofluorescence staining, cells were seeded on coverslips. On the next day, the cells were fixed with 4% methanol-free formaldehyde (Thermo Fisher Scientific; PI28906), and subsequently permeabilized with 0.5% Triton X-100 in PBS. Blocking was performed with 10% FBS + 0.5% Triton X-100 in PBS. Primary antibody incubation was performed for 1 h in a wet chamber. The following primary antibodies were used at a 1:500 dilution in blocking solution: a homemade SAMD1 antibody recognizing the SAM domain, an HA-antibody (Merck; 11867423), and an N-cadherin antibody (Thermo Fisher Scientific; 33-3900). Next, the cells were washed three times with 0.5% Triton X-100 in PBS. Secondary antibody incubation was conducted using Alexa Fluor 488 and 546 coupled antibodies (Thermo Fisher Scientific; A-11008, A-11081; A-11001) at a 1:1000 dilution. To stain the actin cytoskeleton, cells were stained with 1x Phalloidin-California Red Conjugate (Santa Cruz; sc-499440) for 20 min. Following three washing steps, the coverslips were mounted onto microscopy slides using ProLong™ Diamond mounting medium (Thermo Fisher Scientific; P36961). Photos were taken using a Leica DM 5500 microscope.

### Proliferation Assay

To determine proliferation rates, cells were seeded in technical triplicates on 6-well plates at a density of 5×10^4^ cells per well. Cell viability was determined 1, 3, and 7 days after seeding for BxPC3 cells and 1, 3, and 5 days after seeding for PaTu8988t cells using the MTT assay. Therefore, 90 μl of 5 mg/ml thiazolyl blue ≥98% (Carl Roth; 4022) was added to each well. After 1 h, the medium was aspirated, and stained cells were dissolved in 400 μl of lysis buffer (80% isopropanol, 10% 1 M HCl, 10% Triton X-100) and diluted further with PBS if necessary. Absorption was measured at 595 nm using a plate reader. All values were normalized to day 1 to compensate for variations in seeding density. The mean value of three biological replicates was determined.

### Wound Healing Assay

To determine the migration rate of SAMD1 knockout cells, PaTu8988t and BxPC3 cells were seeded in culture inserts (Ibidi; 80209). A total of 70 µl of cell suspension at a density of 6×10^5^ cells per ml was applied. On the next day, the insert was directly removed for BxPC3 cells whereas PaTu8988t cells were starved with medium containing 0.5% FBS for 6 h before removing the insert. Photos were taken using an Olympus CKX53 microscope. After 7 h for BxPC3 cells and after 24 h for PaTu8988t cells, photos were taken on the same spots and the cell-free area was measured for both timepoints using ImageJ Fiji (version: 2.1.0/1.53r).

### Transwell Migration Assay

To determine whether SAMD1 KO has any effects on the migratory potential of PaTu8988t cells, transwell migration assays were performed. Therefore, transwell inserts with a pore size of 8.0 µm (BD Biosciences; 353097) were placed in the wells of a corresponding 24-well plate (Corning; 353504) containing 600 µl serum-free DMEM, high glucose, and GlutaMAX™ Supplement with or without 5% FBS as a chemoattractant. 2×10^4^ PaTu8988t cells in 300 µl serum-free DMEM medium were seeded per transwell insert. The cells were allowed to migrate through the filter for 18 h. Non-migrated cells were removed from the upper transwell insert by wiping them out and performing thorough washing steps in PBS. The migrated cells present on the bottom side of the transwell filter were fixed in methanol for at least 3 minutes and stained with crystal violet solution (0.2% in 20% methanol, 1:5 dilution in dH_2_O) for 10 minutes at room temperature. Membranes were washed in aqua bidest and dried prior to fixing them on microscopy coverslips using Vectashield® with DAPI (Vector Laboratories; H-1200). Evaluation of migrated tumor cells was performed under a Leica DMI3000B microscope. Migrated cells were counted in seven visual fields per filter using ImageJ (version: 2.0.0-rc-43/1.52n). Migration was depicted relative to control.

### Time Lapse Analysis

To perform time lapse analysis, PaTu8988t cells were seeded on collagen-coated 6-well plates at a density of 5×10^4^ cells per well. Coating was performed with 30 mg collagen (Merck; 50201) per ml acetic acid (0.02 M) for 2 h at 37 °C. Afterwards, the plates were washed three times with PBS before seeding the cells.

On the next day, the cells were placed in a Zeiss LSM 780 microscope and every 10 min photos were taken for 24 h using a 10x DIC objective. Migration of cells was analyzed using the “Time Lapse Analyzer” (University of Ulm, version tla_src_v01_33). As a setup file, “DIC tracking 1” was used. Migration was measured in µm per min.

### Measurement of cell shape

To determine the cell shape, cells were seeded on 6-well plates at low density (3×10^4^/well). Photos were taken three days after seeding. For treatment with ADH-1, cells were seeded on 24-well plates (1×10^4^) and treated directly after seeding. Photos were taken one day later using an Olympus CKX53 microscope. For each condition, three photos were taken and 10 cells per photo were analyzed by measuring the circularity of single cells using ImageJ Fiji (version: 2.1.0/1.53r).

### RNA Preparation

For RNA isolation, cells were cultivated on 6-well plates up to 80-100% confluency. RNA was prepared according to the manufacturer’s manual using the RNeasy Mini Kit (Qiagen; 74004) including an on-column DNA digest.

### cDNA Synthesis

The Tetro cDNA Synthesis Kit (Bioline; BIO-151 65043) was used to transcribe mRNA into cDNA according to the manufacturer’s manual. Samples were incubated at 45 °C for 50 min followed by 5 min at 85 °C to inactivate Tetro RT. Subsequently, cDNA was diluted 1:20 for use in RT-qPCR.

### RT-qPCR

For analysis by real-time quantitative PCR, MyTaq™ Mix (Bioline; BIO-25041) was used. For gene expression analysis, values were normalized to GAPDH. Primers are displayed in **Supplementary Table S2.**

### Ectopic Co-immunoprecipitation

All ectopic coimmunoprecipitation (Co-IP) experiments were performed in HEK293 cells. Cells were seeded in 10-cm dishes at 2×10^6^ cells per dish. One day later, the expression constructs for 3xHA or FLAG–tagged proteins were transfected using Polyethylenimine, Linear, MW 25000, Transfection Grade (Polysciences; 23966) and Opti-MEM™ (Thermo Fisher Scientific; 31985062). When the interaction of two proteins should be studied in the presence of a KDM1A inhibitor, the medium was exchanged 5 h after transfection to medium containing either DMSO or 20 nM ORY-1001 (Cay19136; Biomol).

Two days after transfection, extract was prepared using Co-IP buffer (50 mM Tris/Cl (pH=7.5), 150 mM NaCl, 1% Triton X-100, 1 mM EDTA, 10% glycerol, 1xPIC (cOmplete™, Protease Inhibitor Cocktail (Roche; 04693116001), 0.5 mM PMSF). ANTI-FLAG M2 Affinity Gel (Merck, A2220) beads were equilibrated by washing two times with 1× TBS and one time with Co-IP buffer. To bind FLAG-tagged proteins, extracts were added to 50 µl of beads and incubated for approximately 3 hours head over tail at 4°C. After incubation, three washing steps with Co-IP buffer were performed. The FLAG beads were boiled for 3 min in 2× Laemmli buffer without L-mercaptoethanol. Subsequently, L-mercaptoethanol was added to the supernatant and the samples were analyzed via western blotting.

For Co-IP experiments, the following constructs were used:

#### SAMD1

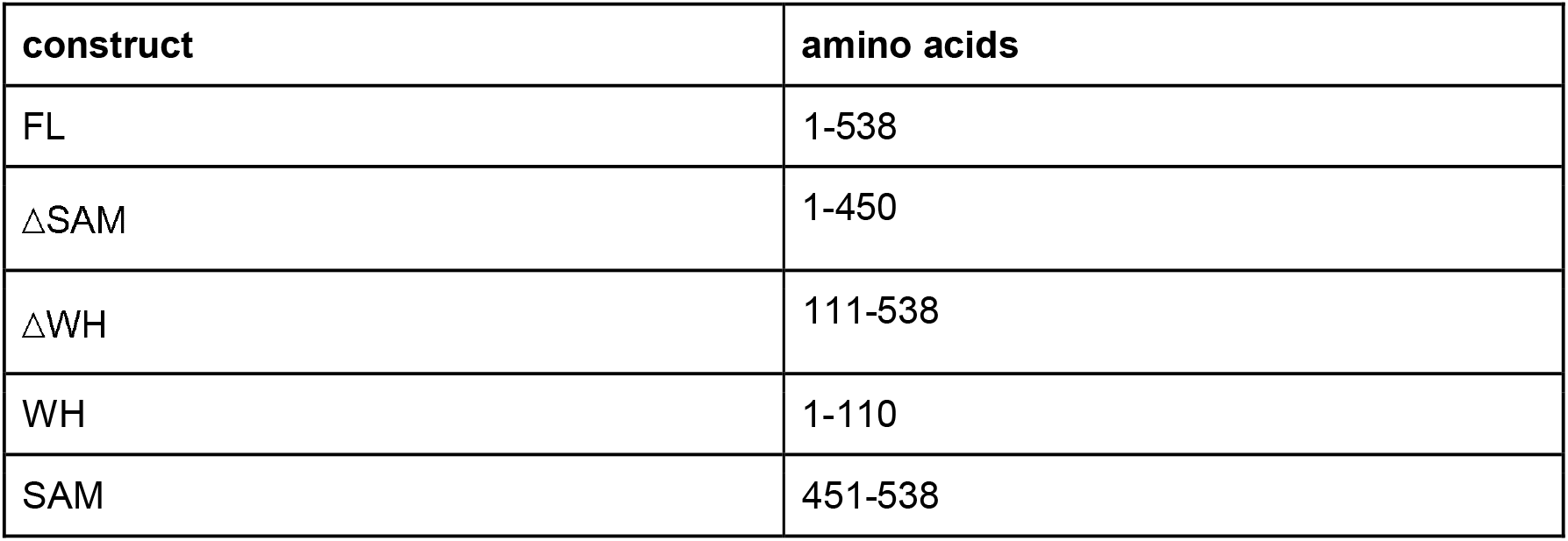

#### KDM1A

KDM1A:
| construct | amino acids |
| --- | --- |
| FL | 1-852 |
| IDR/SWIRM/AOD-N | 1-417 |
| Tower | 418-513 |
| AOD-C | 514-852 |
| IDR | 1-172 |
| SWIRM | 173-272 |
| AOD-N | 273-417 |

### Ubiquitination Assay

To perform a ubiquitination assay, HEK 293 cells stably overexpressing 3xHA-tagged ubiquitin were seeded on 15 cm dishes and transfected one day afterwards with the respective FLAG-tagged constructs using Polyethylenimine, Linear, MW 25000, Transfection Grade (Polysciences; 23966) and Opti-MEM™ (Thermo Fisher Scientific; 31985062). Before preparing extracts, cells were treated with 10 µM MG-132 (M7449; Merck) or DMSO as control for 5 h. Extract preparation and IP were performed according to the ectopic co-immunoprecipitation protocol (see above) and samples were analyzed via western blotting.

### Histone Demethylase Assay

The protocol for a histone demethylase assay was modified after Laurent et al (2015). Wild-type HEK293 cells, HEK293 cells stably overexpressing FLAG-KDM1A and HEK293 cells with SAMD1 KO stably overexpressing FLAG-KDM1A were used. Per reaction, an extract was prepared with buffer A (10 mM HEPES (pH=7.6), 3 mM MgCl_2_, 300 mM KCl, 5% glycerol, 0.5% NP-40, 1x PhosSTOP™ (Roche; 4906845001), 0.5 mM PMSF, 1 µg/µl pepstatin, 10 µg/µl aprotinin, 10 µg/µl leupeptin) using five 15 cm dishes.Subsequently, the extract was diluted by half with buffer B (10 mM HEPES (pH=7.6), 3 mM MgCl_2_, 10 mM KCl, 5% Glycerol, 0.5% NP-40, 1x PhosSTOP™ (Roche; 4906845001), 0.5 mM PMSF, 1µg/µl Pepstatin I 10 µg/µl Aprotinin I 10 µg/µl Leupeptin). 50 µl of ANTI-FLAG M2 Affinity Gel (Merck, A2220) beads were equilibrated with buffer B. Extracts were added to the equilibrated beads and incubated for 3 h head over tail at 4 °C. Next, the beads were washed three times with buffer B and two volumes (100 µl) of demethylase buffer (50 mM Tris/Cl (pH=8.5), 50 mM KCl, 5 mM MgCl_2_, 0.5% BSA, 5% glycerol, 0.5 mM PMSF, 1 µg/µl pepstatin, 10 µg/µl aprotinin, 10 µg/µl leupeptin, 500 μg/mL FLAG® peptide (Merck; F3290)) were added to the beads. To start the demethylase reaction, 3 µg of calf histones (Merck; H9250) was added, and the samples were incubated for 4 h at 37 °C while shaking. The reaction was stopped by boiling the samples for 5 min in 5x lämmli buffer. Subsequently, the samples were analyzed via western blotting.

### Extract preparation and IP for Mass Spectrometry

For IP mass spectrometry, PaTu8988t cells with SAMD1 KO stably expressing either FH-ER as a control or FH-ER SAMD1 were used. For each construct, 20 15 cm dishes were seeded and the nuclear translocation of SAMD1 was induced 24 h before extract preparation by adding 200 nM 4-OHT (Merck; 68392-35-8). After collection, cells were centrifuged at 2,000 rpm and 4 °C for 10 min. The cell pellet was resuspended in 5x pellet volume hypotonic buffer (10 mM Tris (pH=7.3), 10 mM KCl, 1.5 mM MgCl_2_, 0.2 mM PMSF, 10 mM ß-mercaptoethanol, 1xPIC (cOmplete™, Protease Inhibitor Cocktail (Roche; 04693116001)) and shaken at 4 °C for 10-15 min. Next, cells were centrifuged at 2,500 rpm and 4 °C for 10 min. The cell pellet was resuspended again in 5x pellet volume hypotonic buffer and denounced 40x in a cell douncer. To remove cell debris, lysates were centrifuged at 3,500 rpm and 4 °C for 15 min. The pellet was resuspended in 1x pellet volume low salt buffer (20 mM Tris/Cl (pH=7.3), 20 mM KCl, 1.5 mM MgCl_2_, 0.2 mM EDTA, 25% glycerol, 0.2 mM PMSF, 10 mM ß-mercaptoethanol, 1xPIC) and dounced 10x. The sample was shaken in a thermomixer and 0.66x pellet volume of high salt buffer (20 mM Tris/Cl (pH=7.3), 1.2 M KCl, 1.5 mM MgCl_2_, 0.2 mM EDTA, 25% glycerol, 0.2 mM PMSF, 10 mM β-mercaptoethanol) was added dropwise. The extract was shaken for 45 min and centrifuged afterwards for 30 min at 13,000 rpm and 4 °C.

The supernatant containing the proteins was transferred to a a Slide-A-Lyzer™ G2 Dialysis Cassette (3.5K) (Thermo Fisher Scientific; 87724) and dialyzed against 3 L of dialysis buffer (20 mM Tris/Cl (pH=7.3), 100 mM KCl, 0.2 mM EDTA, 20% glycerol, 0.2 mM PMSF, 1 mM DTT) overnight.

To perform the FLAG-IP, the material was retrieved from the dialysis chambers and centrifuged at 13,000 rpm and 4 °C for 30 min. Afterwards, a benzonase® nuclease (Merck;70664) digest (1 µl of benzonase per 500 µl extract) was performed for 1 h on ice. 40 µl of ANTI-FLAG M2 Affinity Gel (Merck, A2220) beads were equilibrated per IP by washing once with TAP buffer (50 mM Tris/Cl (pH=7.9), 100 mM KCl, 5 mM MgCl_2_, 0.2 mM EDTA, 10% glycerol, 0.1% NP-40, 0.2 mM PMSF, 1 mM DTT), three times with 100 mM glycine (pH=2.5), once with 1 M Tris/Cl (pH=7.9) and finally once again with TAP buffer. Subsequently, the extracts were added to the prepared beads and incubated for 3 h, head over tail at 4 °C. Afterwards the beads were washed 3x with TAP buffer and 3x with 50 mM ammonium hydrogen carbonate. The washed beads were then sent in for mass-spectrometry analysis at the Biomedical Center Munich, protein analysis unit (Head: Axel Imhof), where the enriched proteins were analyzed using a Q Exactive HF Orbitrap Mass Spectrometer, as described previously (Weber et al., 2023).

### Endogenous Co-IP

For endogenous Co-IP between SAMD1 and KDM1A, an extract was prepared according to the extract preparation protocol for mass spectrometry and the extract was dialyzed overnight (see above). For each IP, one 15 cm dish of PaTu8988t cells was used. Dynabeads™ Protein A (Thermo Fisher Scientific; 10008D) were equilibrated with TAP buffer (50 mM Tris/Cl (pH=7.9), 100 mM KCl, 5 mM MgCl_2_, 0.2 mM EDTA, 10% glycerol, 0.1% NP-40, 0.2 mM PMSF, 1 mM DTT) and subsequently the extract was precleared for 30 min with 10 µl of beads per IP before adding 2 µg of antibody per IP. Self-made IgG and SAMD1 antibodies and an anti-KDM1A (Abcam; AB17721) antibody were applied. After incubation for 3h, 20 µl of equilibrated Dynabeads™ Protein A per IP was added and incubated for another 2 h. The beads were washed 3x with TAP buffer before boiling in 2x Lämmli buffer.

### Chromatin Preparation

To prepare chromatin, cells were seeded on 15 cm plates at 3×10^6^ cells per plate and cultivated until reaching 70-90% confluency. First, 1% formaldehyde was added to the medium and the plates were slowly swayed for 10 min to fix the cells. The fixation was stopped by adding 125 mM glycine for 5 min. Subsequently, the cells were washed twice with PBS and scraped in 1 ml cold buffer B (10 mM HEPES/KOH (pH=6.5), 10 mM EDTA, 0.5 mM EGTA, 0.25% Triton X-100) per 15 cm plate. All plates containing the same cell line were pooled in a 15 ml tube. The tubes were centrifuged for 5 min at 2000 rpm and 4 °C. The supernatant was removed, and the pellet was resuspended in 1 ml cold buffer C (10 mM HEPES/KOH (pH=6.5), 10 mM EDTA, 0.5 mM EGTA, 200 mM NaCl) per 15 cm plate followed by a 15 min incubation time on ice. Then the tubes were centrifuged with the same settings as mentioned before. After removing the supernatant, the pellet was resuspended in 200 µl cold buffer D (50 mM Tris/HCl (pH=8.0), 10 mM EDTA, 1% SDS, 1xPIC (cOmplete™, Protease Inhibitor Cocktail (Roche; 04693116001)) per 15 cm plate, vortexed, and incubated for 10-20 min on ice. For shearing the chromatin, the samples were sonicated two times for 7 min each using a precooled Bioruptor® (Diagenode). The samples were centrifuged for 10 min at 13.000 rpm and 4 °C. The supernatant contained the sheared chromatin.

### Chromatin Immunoprecipitation

Chromatin-immunoprecipitation (ChIP) for ChIP-qPCR was performed according to the One-day ChIP kit protocol (Diagenode; C01010080). Beads were exchanged to Dynabeads™ Protein A (Thermo Fisher Scientific; 10008D), and Chelex to Chelex 100 Resin (BioRad;142-1253) and ChIP buffer was replaced by a homemade buffer (50 mM Tris/Cl (pH=7.5), 150 mM NaCl, 5 mM EDT, 1% Triton X-100, 0.5% NP-40). For each ChIP, 3 µg of either IgG control antibody (Diagenode; C15410206) or of a specific antibody were applied. For histone marks only 1 µg of antibody was used. The following antibodies were used: a self-made SAMD1 antibody recognizing the SAM domain, a self-made L3MBTL3 antibody recognizing the SAM domain, anti-KDM1A (Abcam; AB17721), anti-H3K3me2 (Diagenode; C15410035), and anti-H3K4me3 (Diagenode; C15410003).

To prepare samples for ChIP-sequencing, the One-day ChIP kit protocol was used as described above, but the DNA-purification was modified. For DNA elution, beads were incubated with 230 µl elution buffer (100 mM NaHCO_3_, 1% SDS) for 30 min at room temperature while shaking. Afterward, the samples were centrifuged at 13.000 rpm for 1 min and 200 µl of supernatant was transferred to a fresh tube. The input DNA was dissolved in 50 µl of dH2O and 150 µl of elution buffer was added to obtain an equal volume in all samples. 8 µl of 5 M NaCl were added to each sample and the samples were incubated at 65 °C overnight to reverse the cross-linking.

On the next day, 8 µL of 1M Tris/Cl (pH=6.5), 4 µL 0.5 M EDTA, and 2 µL of Proteinase K (10 µg/µL) were added to each sample and all samples were incubated at 45 °C for 1 h whilst shaking. DNA was purified using the QIAquick PCR Purification Kit (Qiagen; 28104) whereby all samples prepared with the same antibody were pooled on the same column. To elute the DNA, columns were incubated for 1 min with 30 µl of sterile 2 mM Tris/Cl (pH=8.5), and centrifuged at 13.000 rpm for 1 min.

The concentration of the samples was determined using the Quant-iT™ dsDNA Assay Kit (Thermo Fisher Scientific; Q33120) and the NanoDrop™ 3300 (Thermo Fisher Scientific). At least 4 ng of DNA was used for library preparation.

### Library Preparation and Next Generation Sequencing

Next generation sequencing was performed at the Genomics Core Facility Marburg (Center for Tumor Biology and Immunology, Hans-Meerwein-Str. 3, 35043 Marburg, Germany). For ChIP-Seq, the Microplex library preparation kit v2 (Diagenode, C05010012) was used for indexed sequencing library preparation with chromatin immunoprecipitated DNA. Libraries were purified on AMPure magnetic beads (Beckman; A6388). RNA was prepared as described in *RNA preparation* and integrity was assessed on an Experion StdSens RNA Chip (Bio-Rad; 7007103). RNA-Seq libraries were prepared using the TruSeq Stranded mRNA Library Prep kit (Illumina, 2002059). RNA-Seq and ChIP-Seq libraries were quantified on a Bioanalyzer (Agilent Technologies). Next-generation sequencing was performed on an Illumina NextSeq 550.

### Bioinformatic analyses

ChIP-Seq data were mapped to the human genome hg38 using bowtie (Langmead et al., 2009), allowing 1 mismatch. BigWig files were obtained using DeepTools/bamCoverage (Ramirez et al., 2014). Significant peaks were obtained using Galaxy/MACS2 (2.2.7.1) (Zhang et al., 2008). Heatmaps and profiles were created using Galaxy/DeepTools (Ramirez et al., 2014). The top target genes were identified based on the SAMD1 ChIP-Seq signal at promoters.

RNA-Seq data were aligned to the human transcriptome (GenCode 43) using Galaxy/RNA-Star (2.7.10b) (Dobin et al., 2013). Differentially expressed genes were obtained using DeSeq2 (2.11.40.7) (Love et al., 2014). Genes with a log2-fold change of more than 0.5 and a p-value lower than 0.01 were considered significantly dysregulated. Gene set enrichment analysis was performed using GSEA software with standard settings (Subramanian et al., 2005).

The following internet databases and tools were used: Galaxy Europa (Galaxy Community, 2022), DeepTools (Ramirez et al., 2014), GREAT (4.0.4) (McLean et al., 2010), Bioconductor/R (Gentleman et al., 2004), GSEA (4.3.2) (Subramanian et al., 2005), GePIA (Tang et al., 2017), GDC Data Portal (Weinstein et al., 2013) and Kaplan-Meier-Plotter (Nagy et al., 2021).

The following public ChIP-Seq data were used: ChIP-Seq of H3K4me3 (GSM945261)(Thurman et al., 2012) and RNA Polymerase II (GSM1010788)(Gertz et al., 2013) in PANC-1 cells.

### Statistical analysis

Statistical analysis was performed as described in the figure legends. Error bars indicate standard deviation (SD). The significance of the qPCR results was analyzed via ANOVA or Student’s t-tests. The significance of the GSEA was evaluated by the GSEA software. The significance of changes in SAMD1 ChIP-Seq levels was evaluated using a two-sided Kolmogorov-Smirnov test. All biological experiments were performed in at least three replicates. RNA-Seq was performed with three replicates, using three independent SAMD1 KO clones.

**Supplementary Videos**

Time Lapse videos of migration of PaTu8988t Control (Video 1) and SAMD1 KO cells (Video 2) for 24h. Every 10 min photos were taken.

---

**Supplementary Figure 1:**
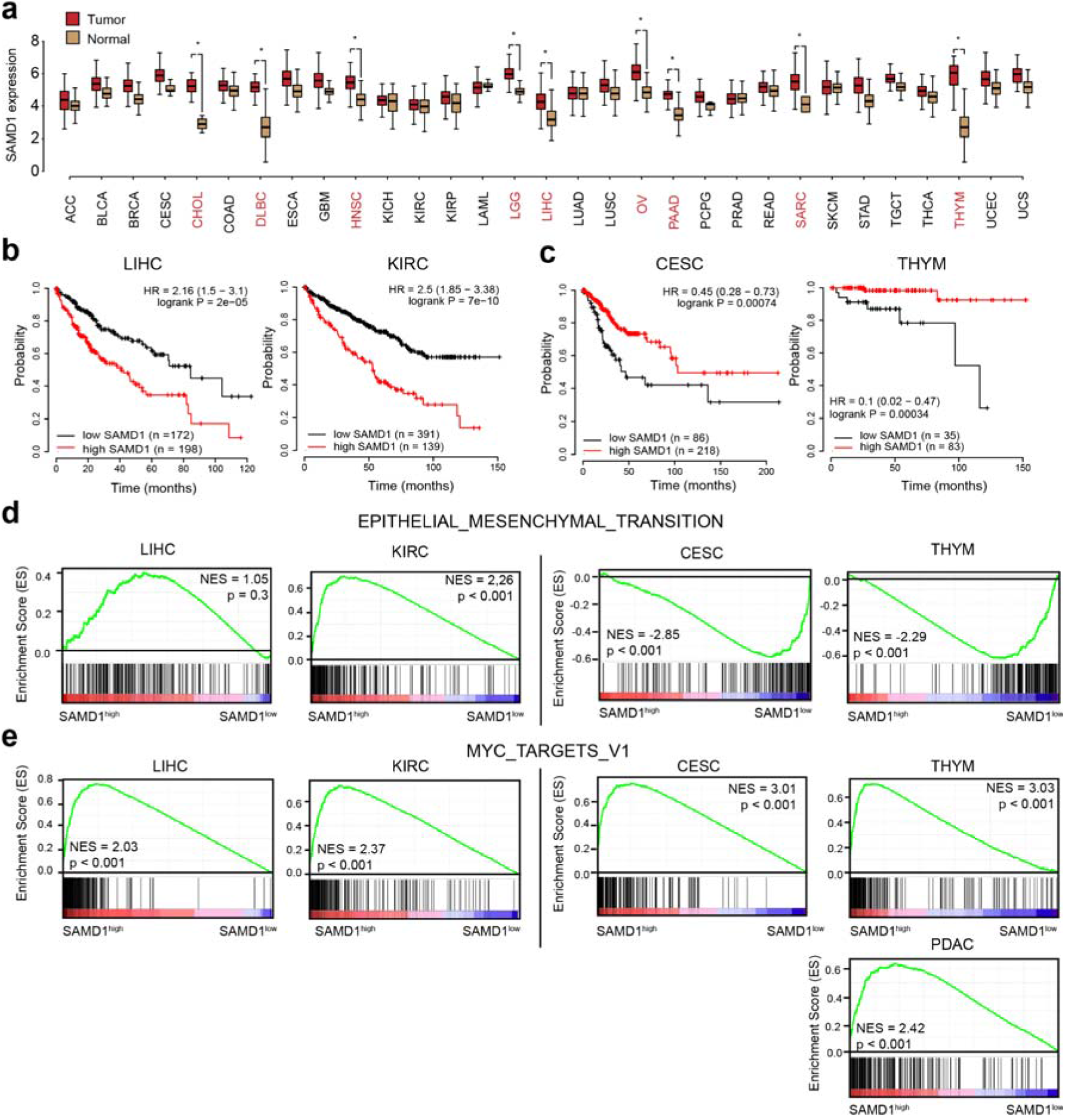
SAMD1 has distinct roles in various cancer types. a) Expression of *SAMD1* in cancer versus normal tissues. Data from TCGA (Weinstein et al., 2013) and visualized via GePIA (Tang et al., 2017). Cancer types in red indicate those with significantly upregulated *SAMD1* expression. b) Kaplan-Meier survival curves (overall survival) in liver hepatocellular carcinoma (LIHC) and kidney renal clear cell carcinoma (KIRC), using auto-selected cut-offs. High SAMD1 expression correlates with a worse prognosis. c) Kaplan-Meier survival curves (overall survival) in cervical cancer (CESC) and thymoma (THYM). High *SAMD1* expression correlates with a better prognosis. Data in b) and c) are derived from TCGA and visualized via the Kaplan-Meier-Plotter tool (Nagy et al., 2021) using auto-selected cut-off. d) GSEA of the epithelial-mesenchymal transition (EMT) pathway in the cancer types presented in b) and c). e) GSEA of the MYC target genes in the cancer types presented in b) and c) and of PDAC. In d) and e), tissue samples with high *SAMD1* expression are compared to samples with low *SAMD1* expression.

**Supplementary Figure 2:**
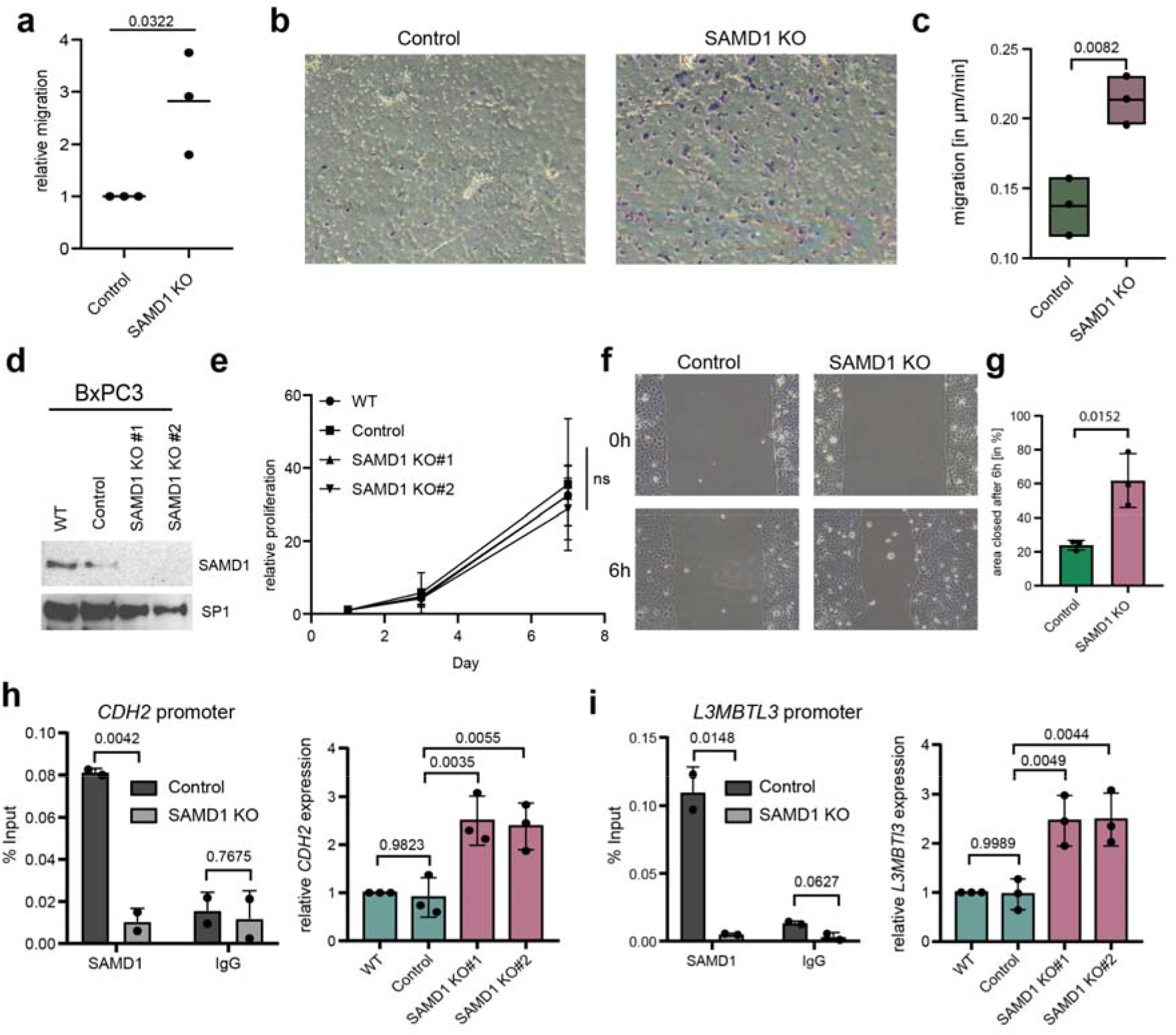
transwell and time-lapse assays in PaTu8988t cells and SAMD1 KO in BxPC3 cells. a) transwell migration assay of PaTu8988t control and SAMD1 KO cells. Data represent the mean ± SD of three biological replicates. Significance was analyzed using Student’s t-test. b) Representative crystal violet staining of one transwell migration assay. c) Migration of PaTu8988t control and SAMD1 KO cells in µm/min based on time-lapse analysis. See also **Supplementary Video 1 and 2**. Data represent the mean ± SD of three biological replicates. Significance was analyzed using Student’s t-test. d) Western blot showing BxPC3 wild-type cells, control cells, and two different SAMD1 knockout clones. e) Proliferation assay of BxPC3 wild-type cells, control cells, and two different SAMD1 knockout clones. Data represent the mean ± SD of three biological replicates. Significance was analyzed using one-way ANOVA. f) Representative picture of one wound healing assay of BxPC3 control cells and one SAMD1 knockout clone. g) Quantification of the wound healing assay from c). Data represent the mean ± SD of three biological replicates, and significance was analyzed using Student’s t-test. h) ChIP-qPCR of *CDH2* promoter in BxPC3 Control and SAMD1 KO cells using IgG or SAMD1 antibodies. Significance was analyzed using Student’s t-test, RT-qPCR showing *CDH2* expression in BxPC3 wild-type cells, control cells, and two different SAMD1 knockout clones. Data represent the mean ± SD of three biological replicates. Significance was analyzed using one-way ANOVA. i) ChIP-qPCR at the *L3MBTL3* promoter in BxPC3 control cells and SAMD1 KO cells, using SAMD1 or IgG antibodies. Significance was analyzed using Student’s t-test, RT-qPCR showing *L3MBTL3* expression in BxPC3 wild-type cells, control cells, and two different SAMD1 knockout clones. Data represent the mean ± SD of three biological replicates. Significance was analyzed using one-way ANOVA.

**Supplementary Figure 3:**
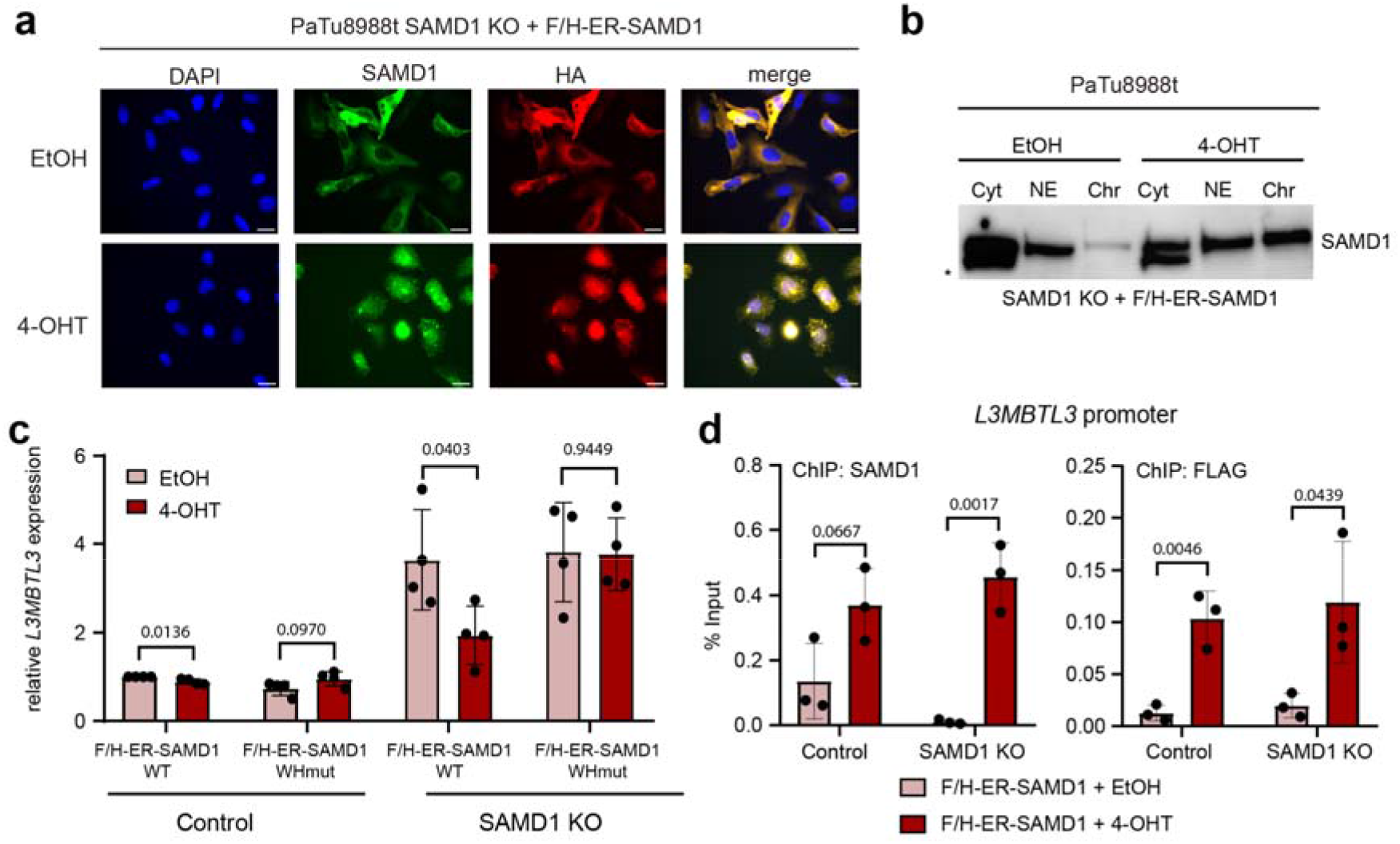
SAMD1 rescue experiments in PaTu8988t cells. a) Immunofluorescence of PaTu8988t SAMD1 knockout cells with or without induction of SAMD1 rescue, Bar=20 µM. b) Western blot after fractionation of PaTu8988t SAMD1 knockout cells with or without induction of SAMD1 rescue. c) RT-qPCR showing *L3MBTL3* expression with or without induction of SAMD1 rescue in PaTu8988t control and SAMD1 KO cells. WHmut=RK-45/46-AA mutation of SAMD1. Data represent the mean ± SD of four biological replicates. Significance was analyzed using Student’s t-test. d) SAMD1 ChIP-qPCR at the *L3MBTL3* promoter with or without induction of SAMD1 rescue in PaTu8988t Control and SAMD1 KO cells. Data represent the mean ± SD of three biological replicates. Significance was analyzed using Student’s t-test.

**Supplementary Figure 4:**
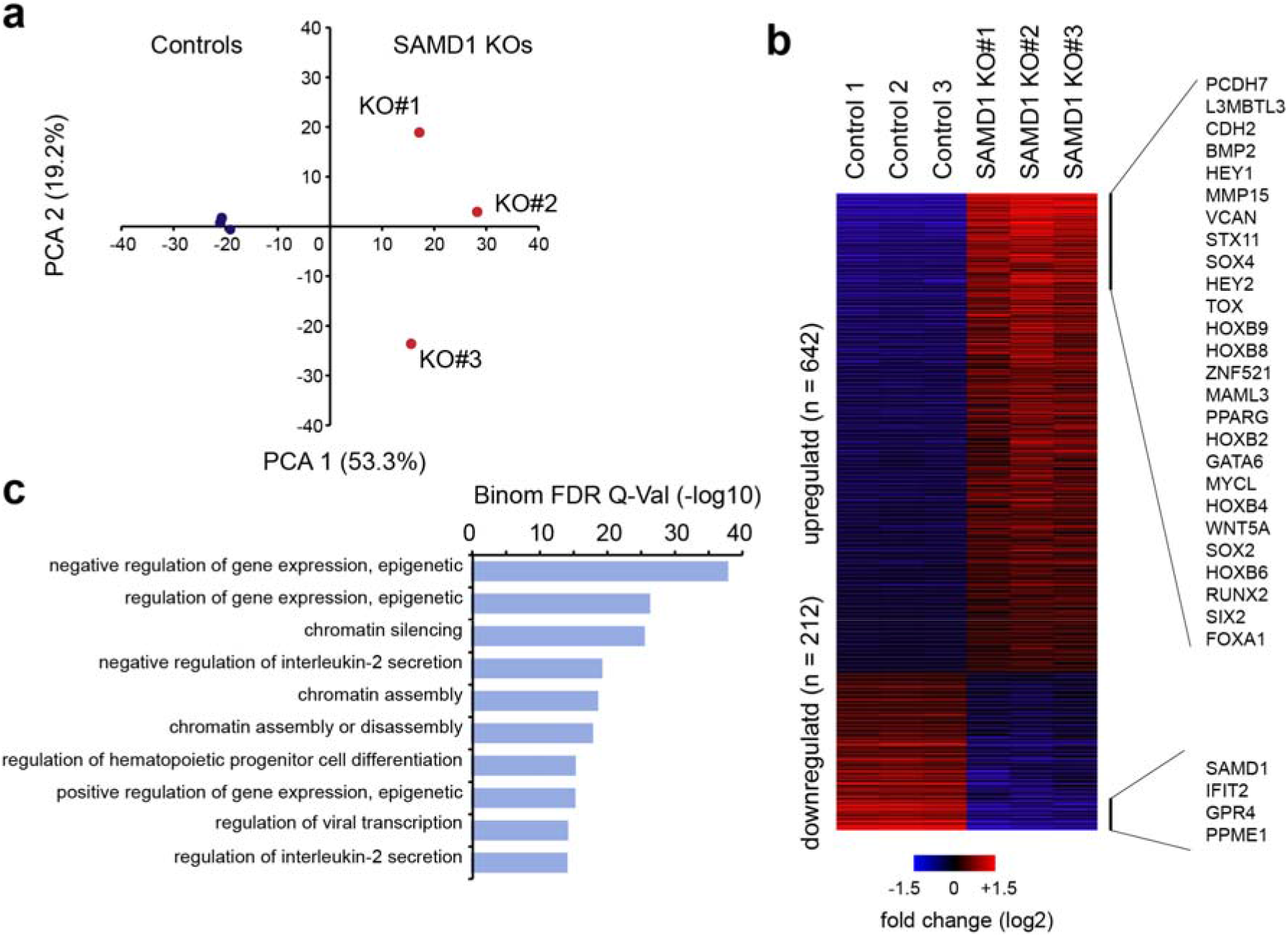
Transcriptional regulation of SAMD1 in PaTu8998t cells. a) Principal component analysis (PCA) of RNA-Seq data upon SAMD1 KO. Three clonally independent SAMD1 KO clones were used. b) Heatmap of the significantly dysregulated genes. Examples of the most dysregulated genes are shown on the right. c) Gene ontology analysis of SAMD1 genomic targets using GREAT (McLean et al., 2010).

**Supplementary Figure 5:**
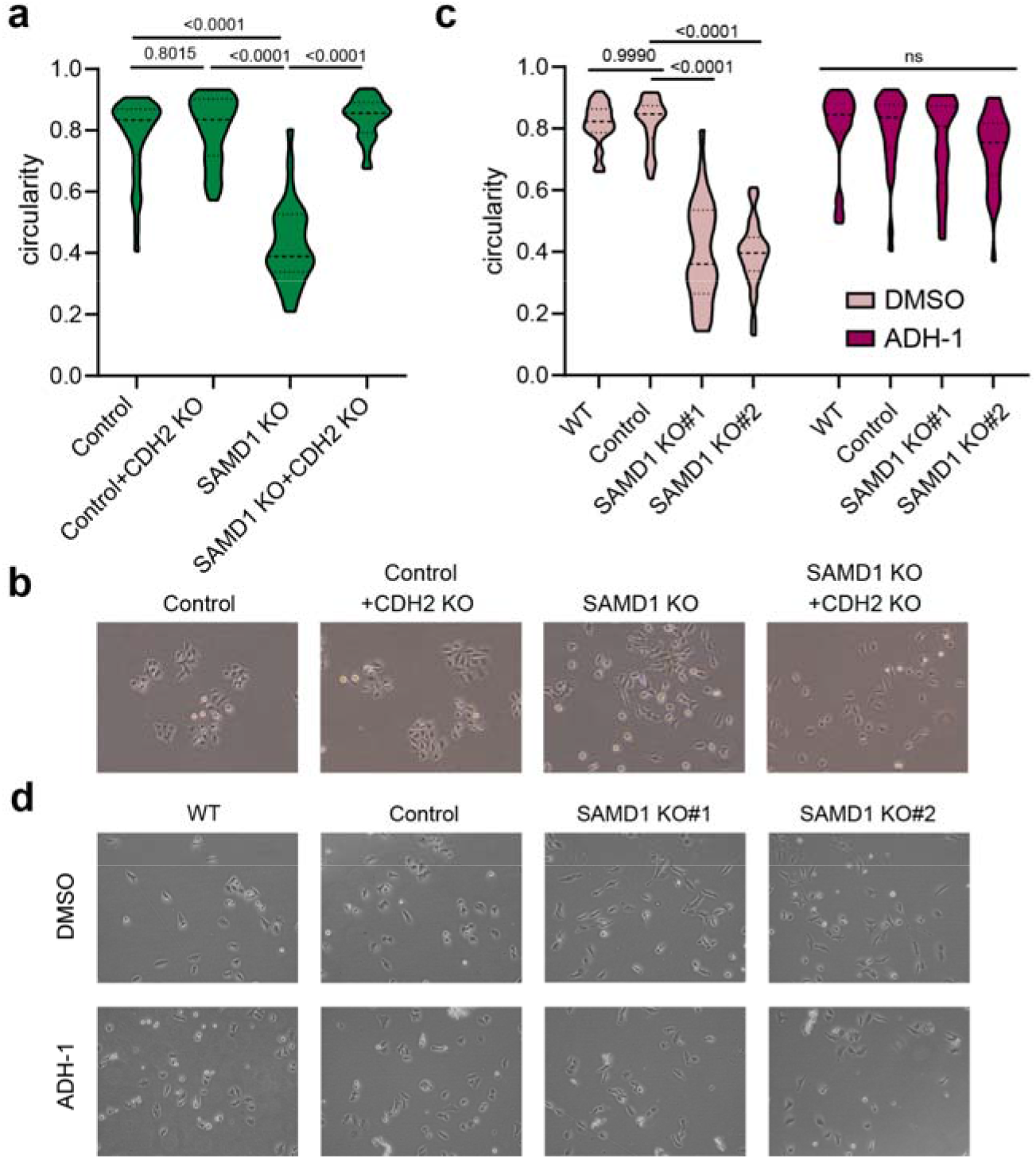
CDH2 KO or inhibition rescues the migration phenotype upon SAMD1 deletion. a) Cell shape of control, CDH2 KO, SAMD1 KO, and CDH2/SAMD1 double KO PaTu8988t cells. Circularity was determined using ImageJ Fiji. Significance was analyzed using one-way ANOVA. b) Example bright field microscopy for a). c) Cell shape of PaTu8988t wild-type cells, control cells and two different SAMD1 knockout clones with or without application of the N-cadherin inhibitor ADH-1. Circularity was determined using ImageJ Fiji. Significance was analyzed using one-way ANOVA. d) Example bright field microscopy for c).

**Supplementary Figure 6:**
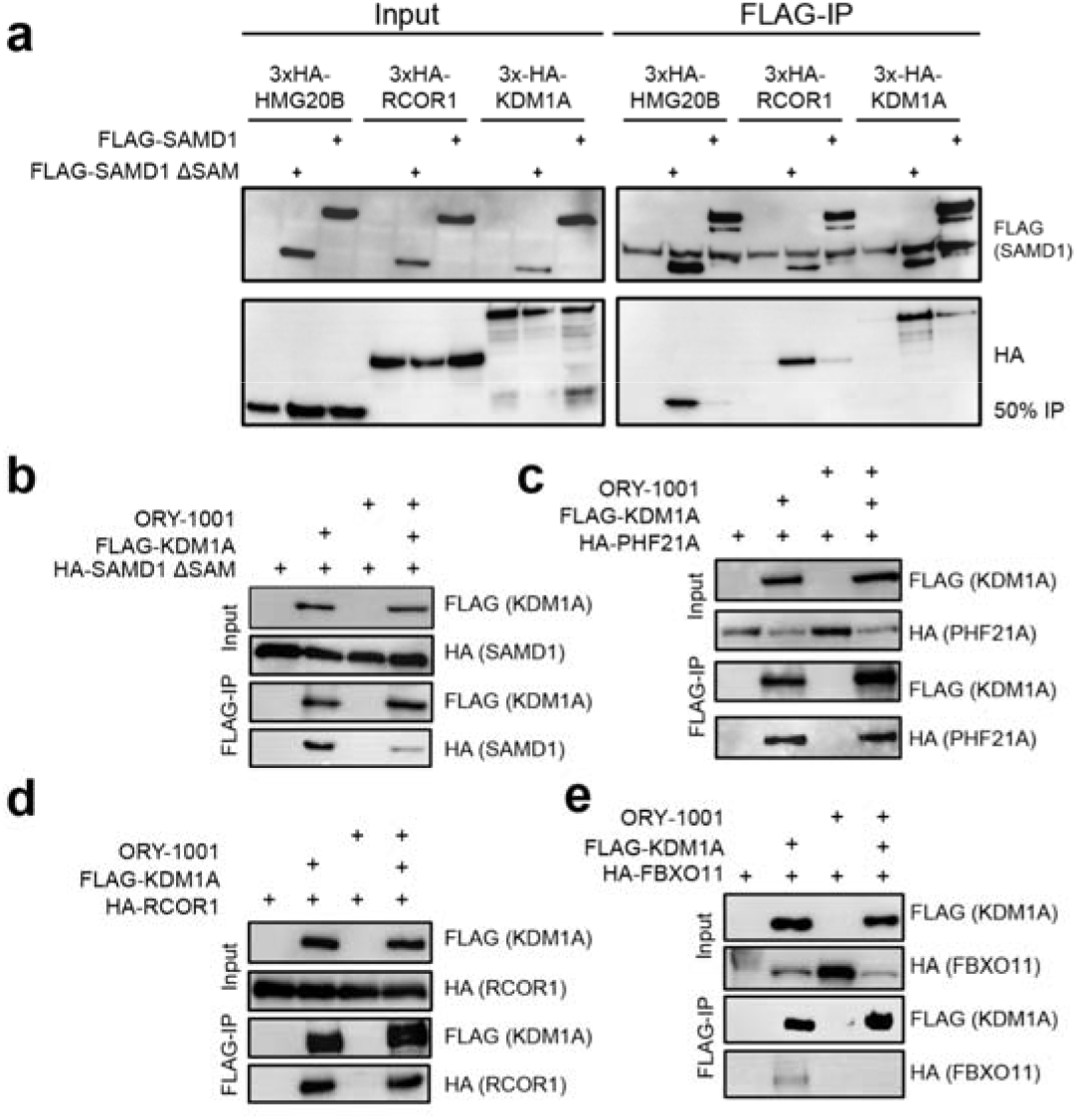
SAMD1/KDM1A interaction is influenced by SAM domain and ORY-1001. a) Co-immunoprecipitation in HEK293 cells showing the interaction between SAMD1_FL or SAMD1ΔSAM and the KDM1A-complex. b) Co-immunoprecipitation in HEK293 cells showing the interaction between SAMD1ΔSAM and KDM1A upon treatment with the KDM1A inhibitor ORY-1001. c) Co-immunoprecipitation in HEK293 cells showing the interaction between PHF21A and KDM1A upon treatment with the KDM1A inhibitor ORY-1001. d) Co-immunoprecipitation in HEK293 cells showing the interaction between RCOR1 and KDM1A upon treatment with the KDM1A inhibitor ORY-1001. e) Co-immunoprecipitation in HEK293 cells showing the interaction between FBXO11 and KDM1A upon treatment with the KDM1A inhibitor ORY-1001.

**Supplementary Figure 7:**
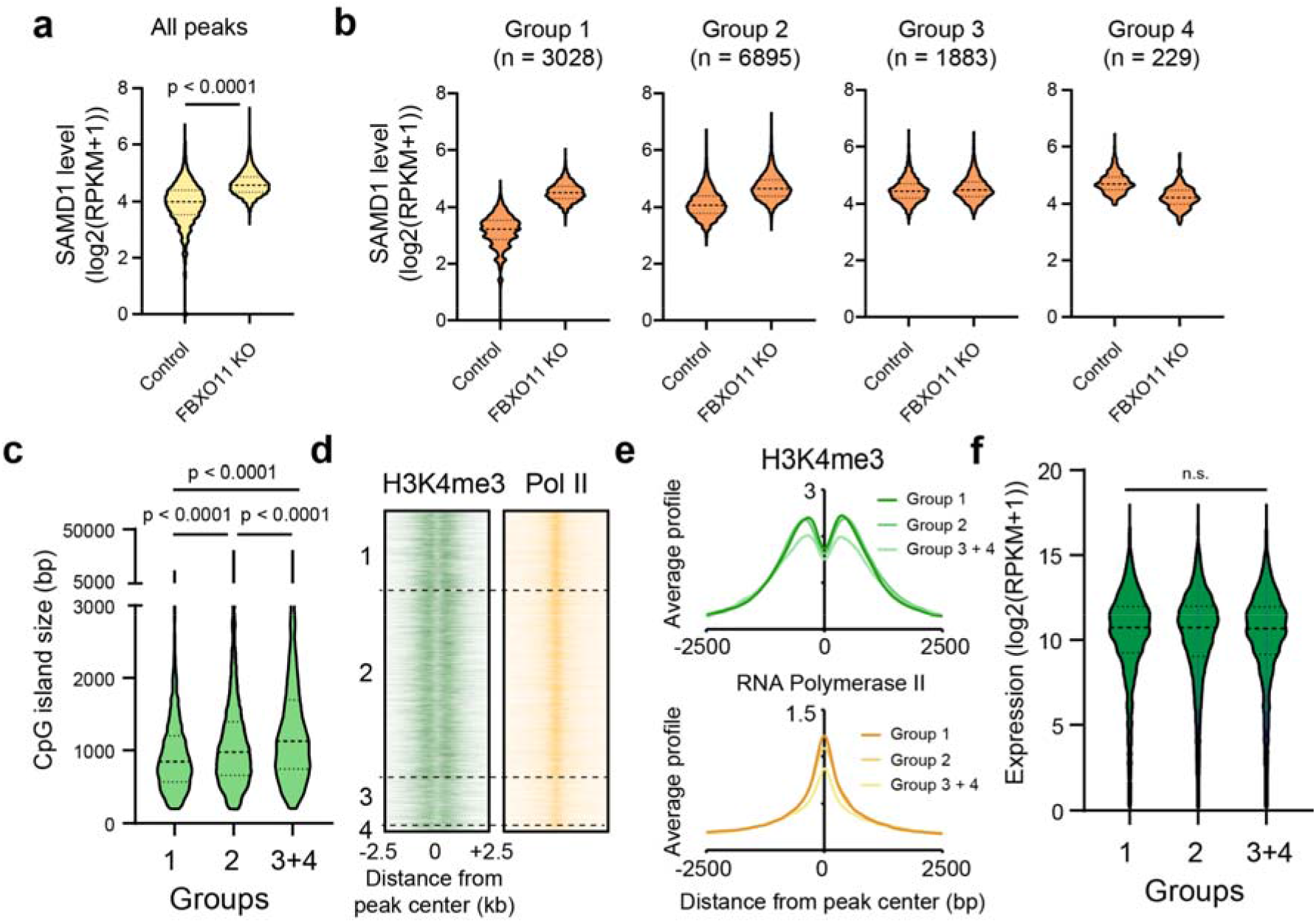
Detailed analysis of the consequence of FBXO11 deletion on SAMD1 chromatin binding. a) Violin plots showing the SAMD1 level in PaTu8988t control and FBXO11 KO cells. Statistical significance was evaluated using a Kolmogorov-Smirnov test. b) SAMD1 levels in four different groups identified in Figure 6g in PaTu8988t control and FBXO11 KO cells. c) CpG island size of four different groups identified in Figure 6c. Statistical significance was evaluated using a Kolmogorov-Smirnov test. d) Heatmap of H3K4me3 and Pol II in the four different groups identified in Figure 6c. e) Profiles of H3K4me3 and RNA Polymer II at the four different groups identified in Figure 6c. f) Expression of genes at the four different groups identified in Figure 6c.

**Table S1.**
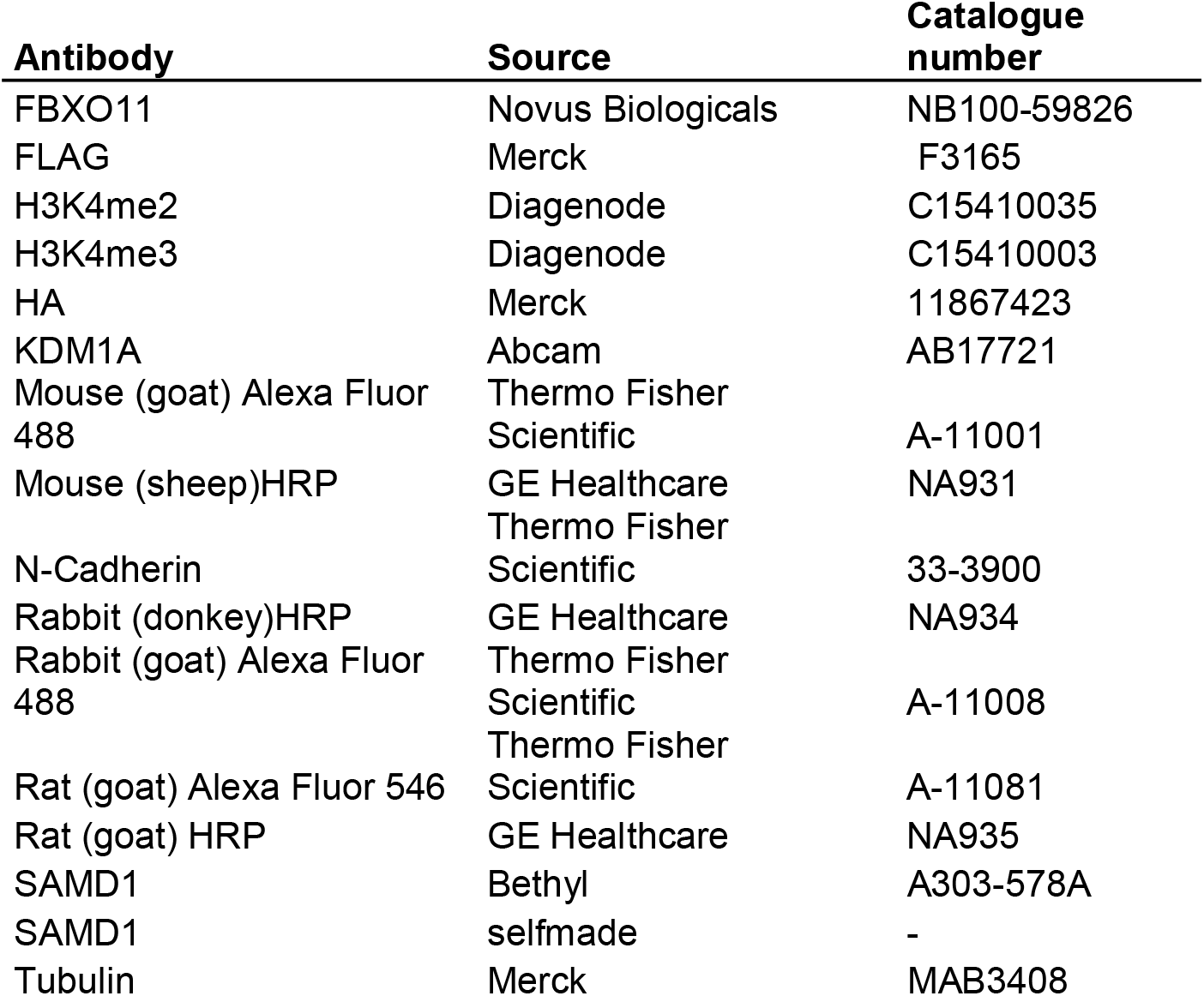
Antibodies.

| **Antibody** | **Source** | **Catalogue number** |
| --- | --- | --- |
| FBXO11 | Novus Biologicals | NB100-59826 |
| FLAG | Merck | F3165 |
| H3K4me2 | Diagenode | C15410035 |
| H3K4me3 | Diagenode | C15410003 |
| HA | Merck | 11867423 |
| KDM1A | Abcam | AB17721 |
| Mouse (goat) Alexa Fluor 488 | Thermo Fisher Scientific | A-11001 |
| Mouse (sheep)HRP | GE Healthcare | NA931 |
| N-Cadherin | Thermo Fisher Scientific | 33-3900 |
| Rabbit (donkey)HRP | GE Healthcare | NA934 |
| Rabbit (goat) Alexa Fluor 488 | Thermo Fisher Scientific | A-11008 |
| Rat (goat) Alexa Fluor 546 | Thermo Fisher Scientific | A-11081 |
| Rat (goat) HRP | GE Healthcare | NA935 |
| SAMD1 | Bethyl | A303-578A |
| SAMD1 | selfmade | - |
| Tubulin | Merck | MAB3408 |

**Table S2.**
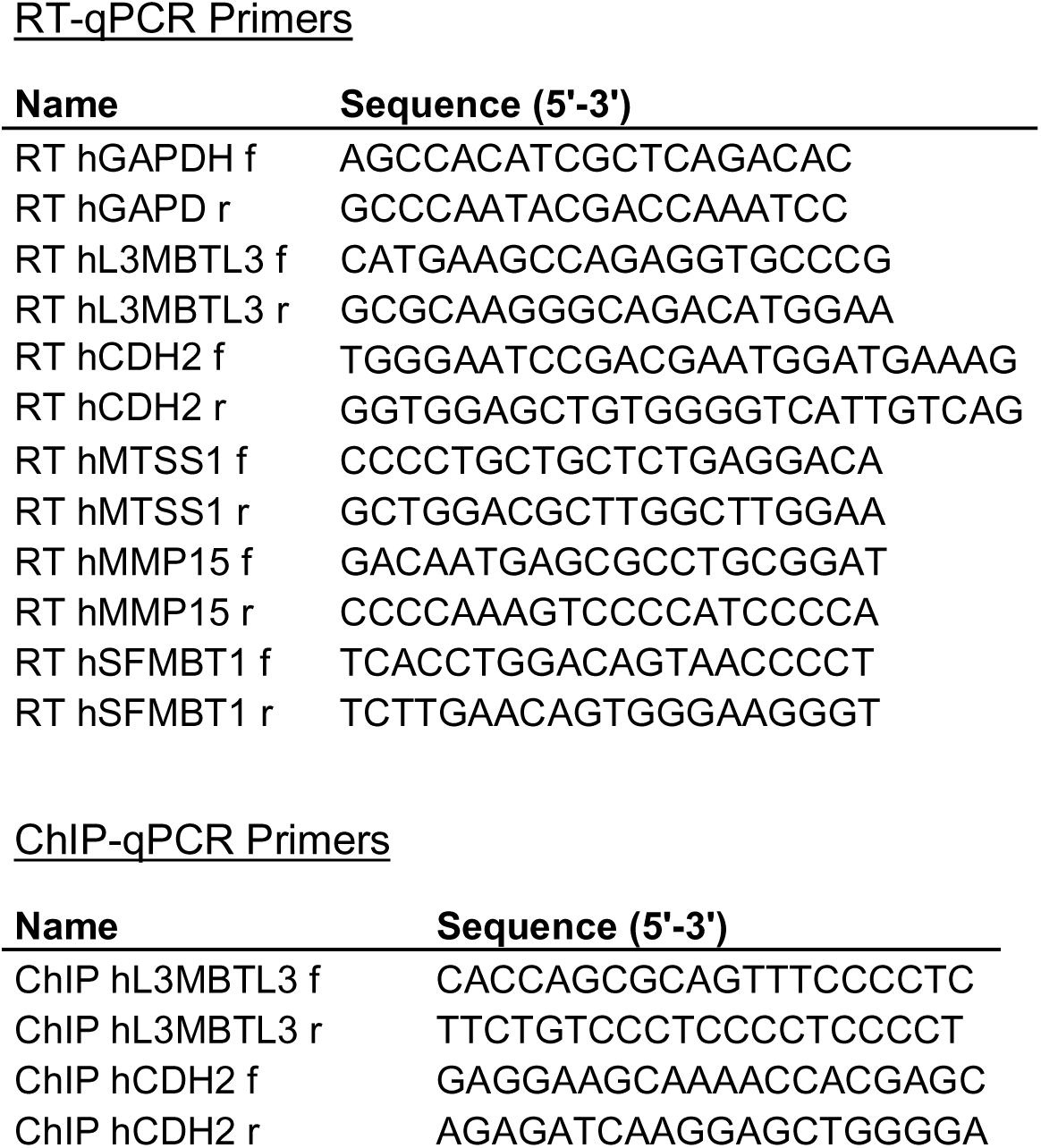
qPCR Primers.

## Notes

### Competing Interest Statement

The authors have declared no competing interest.

